# Origin of Fe ions in ROS production induced in magnetic hyperthermia anti-cancer nanotherapy: release from iron oxide nanoparticles or not?

**DOI:** 10.64898/2026.07.06.736751

**Authors:** Justine Journaux-Duclos, Megi Bejko, Pascal Clerc, Yasmina Al Yaman, Ahmed Abdelhamid, Géraldine Ballon, Corinne Bousquet, Julian Carrey, Stéphane Mornet, Olivier Sandre, Véronique Gigoux

**Affiliations:** Centre de Recherches en Cancérologie de Toulouse, INSERM U1037, Toulouse, France; Institut de Chimie de la Matière Condensée de Bordeaux, Univ. Bordeaux, CNRS, Bordeaux INP, ICMCB, F-33608 Pessac, France; Laboratoire de Chimie des Polymères Organiques, Univ. Bordeaux, CNRS, Bordeaux INP, LCPO, F-33607 Pessac, France; Laboratoire de Physique et Chimie des Nano-Objets, CNRS-UPS-INSA UMR5215, Toulouse, France

**Keywords:** Cancer, magnetic nanoparticles, targeted nanotherapy, magnetic hyperthermia, ROS, cell death

## Abstract

The first and critical reaction in magnetic hyperthermia to induce the death of cancer cells is the production of ROS (reactive oxygen species). We previously showed that it is possible to specifically deliver iron oxide magnetic nanoparticles (IONPs) in the lysosomes of cancer cells and eradicate them by targeted magnetic intra-lysosomal hyperthermia (MILH) *via* the application of a high frequency alternating magnetic field (AMF) without macroscopic temperature elevation. The mechanism involves a local temperature elevation at the IONPs surface which enhances the ROS production through the Fenton reaction; ROS then peroxide the proteins and lipids of the lysosomal membrane, inducing its permeabilization and leading to lysosomal enzymes release and cell death. Fe ions, critical to produce ROS in MILH, were assumed to be released by IONPs. We thus developed PEGylated multi-cores IONPs called NanoFlowers (NF@PEG) presenting or not a SiO_2_ shell (NF@SiO_2_ @PEG), the later preventing the Fe^3+^ release from IONPs. NF@PEG released Fe ions and produced ROS production *in vitro*, in acidic medium mimicking lysosome upon AMF exposure, whereas NF@SiO_2_@PEG did not. Surprisingly, both nanoparticles increased the ROS production in cells, induced lysosome permeabilization and cell death, and slowed down the proliferation of cancer cells with the same efficacy, upon AMF application, indicating that MILH was efficient in absence of Fe^3+^ release from IONPs. In contrast, Ferristatin-II, an iron uptake inhibitor, prevented the ROS production and cell death in MILH induced by both IONPs, elucidating the role of endogenous iron cations responsible for the ROS production ROS in MILH to kill cancer cells.

## INTRODUCTION

Iron oxide magnetic nanoparticles (IONPs), which display low toxicity and good biocompatibility in humans, are the subject of renewed interest in the perspective of cancer nanotherapy ^1^. Exposed to a high frequency alternating magnetic field (AMF), IONPs release heat which can be used to develop biological approaches such as the eradication of cancer cells (magnetic hyperthermia), the on-demand release of drugs or the disruption of extracellular matrix of the tumor microenvironment ^2–5^. Magnetic hyperthermia was authorized to treat high grade brain tumors, when combined with radiotherapy ^6^. However, survival rate is still modest (average gain of 2 months lifetime) and neither radiotherapy nor magnetic hyperthermia therapy can discriminate between normal and cancerous tissues. Moreover, the lack of tissue discrimination as well as the large injected doses to increase magnetic hyperthermia therapy effectiveness, can lead to undesirable side effects ^7^. Currently, magnetic hyperthermia in cancer treatment has obtained a renewed interest with the FDA approval of prostate cancer in combination with radiotherapy, in 2018. In order to overcome the limits imposed by macroscopic magnetic hyperthermia using large doses of magnetic nanoparticles mainly injected intratumorally and causing adverse effects on surrounding healthy tissues, alternating magnetic intra-lysosomal hyperthermia (MILH), also called intracellular magnetic hyperthermia, investigated the local release of thermal energy by targeted IONPs within the lysosome to induce lysosomal death pathways and kill the cancer cells without macroscopic temperature rise ^8–10^. The mechanism involves a local temperature elevation at the IONPs surface which enhances the ROS (reactive oxygen species) production through the Fenton reaction (Fe^2+^+H_2_ O_2_ →Fe^3+^+OH^−^+OH^•^); ROS then peroxide the proteins and lipids of the lysosomal membrane, inducing its permeabilization and leading to lysosomal enzymes release and cell death ^9–12^. Cell death induced by magnetic hyperthermia was also characterized to be related to ferroptosis, as it depended on Fe ions and ROS production by Fenton reaction and affected the expression of ferroptosis-related proteins ^11,13^.

The degradation of IONPs (γ-Fe_2_O_3_ and Fe_3_O_4_) within fluids mimicking lysosomal composition and pH as well as *in situ* has been described, mainly releasing Fe^3+^ and Fe^2+^ respectively ^14–18^. Conceivably, these newly formed Fe ions may affect the intracellular oxido-reduction reactions and homeostasis of ROS inside cells. Indeed, Fe is an important redox-active transition metal involving production of highly reactive free radicals (•OH and O_2_^•−^) *via* Fenton reaction and Haber-Weiss reactions cycle in living systems ^19,20^. These reactions are dependent on Fe concentration, acidic pH (optimal at pH 3) and temperature (optimal at 40-45°C) ^21,22^. Thus, the combination of the local temperature rise at the IONPs surface induced by AMF with the acidic pH of the lysosomal compartment may enhance the IONPs degradation and the release of Fe ions from IONPs, which participate in the cytotoxic Fenton reaction leading to cell death upon AMF exposure ^23^. However, lysosome is a cellular compartment rich in iron, through the extracellular iron delivering to the lysosomes by endocytosis (*via* the transferrin/transferrin receptor system) and the degradation of intracellular materials (ferritin, old and damaged cellular macromolecules and organelles), making the lysosome a master regulator of cellular iron homeostasis and a crucial determinant of cell death by ferroptosis ^24^. Indeed, the increase of intracellular Fe ions and follow-up Fenton reaction, which elevates ROS levels, lead to ferroptosis cell death. Currently, the origin of Fe ions involved in the Fenton reaction (i.e. either exogenous Fe ions issued from IONPs’ degradation or endogenous intracellular Fe) during magnetic hyperthermia remains misunderstood. To test these hypotheses, we developed PEGylated IONPs constituted by a multi-core iron oxide (*γ*-Fe_2_O_3_) nanoparticle called “NanoFlowers” (NF@PEG) presenting or not a SiO_2_ shell (NF@SiO_2_ @PEG), the later preventing Fe^3+^ release from IONPs. We first analyzed the Fe release from both nanoparticles and their capacity to generate ROS *in vitro*. We then investigated whether NF@PEG and NF@SiO_2_@PEG nanoparticles induce differently intracellular ROS production, lysosome permeabilization and affect cell viability. We also performed same studies, in the presence or not of Ferristatin-II, an iron uptake inhibitor that interferes with transferrin-mediated iron delivery by inducing the internalized degradation of transferrin receptor 1 (TfR1) ^25^. This study allows to elucidate the origin of Fe ions responsible for ROS production during magnetic hyperthermia to kill cancer cells.

## RESULTS AND DISCUSSION

### Silica shell prevents Fe release from iron oxide nanoparticles

We developed and characterized iron oxide superparamagnetic multicore nanoparticles, termed nanoflowers, presenting or not a silica shell preventing Fe ion release from the nanoparticle core (Figure S1). The nanoflowers (NF) were synthesized using polyol route^26^. The magnetic core size, measured by TEM, equaled 17.3 ± 2.6 nm (Figure 1A-B and Figures S1 and S2, Table 1). Half of the nanoflowers batch was coated with a sol-gel silica (NF@SiO2) shell; the thickness of silica shell was determined by TEM at 3.1 ± 0.2 nm. Both nanoflowers types, NF and NF@SiO_2_, were then covered with polyethylene glycol (PEG) to ensure their colloidal stability and reduce opsonization ^27^. NF was modified by phosphonate PEGylated ligands bearing amine functions as terminal groups (PO-PEG-2k-NH_2_) in one-step surface reaction; NF@SiO_2_ was multi-step modified by amination surface, grafting of PEG-2k-CHO through reductive amination of OH end groups of grafted PEG, as illustrated in Figure S1 and previously described for SiO_2_ nanoparticles ^28^. The PEG corona of NF@PEG and NF@SiO_2_ @PEG were both situated in brush conformational regime, as determined from the R_F_/D ratio (2.4 and 2.7 respectively, Table 1). The hydrodynamic diameter of NF@PEG and NF@SiO_2_@PEG were respectively equal to 50.3 ± 4.0 and 64.5 ± 6.9 nm in H_2_O, and their polydispersity index PDI 0.25 ± 0.15 and 0.17 ± 0.02, indicating a low nanoparticles aggregation (Table 1). The positive zeta potential value of NF@PEG (+14 ± 2.0 mV) and NF@SiO_2_@PEG (+22.6 ± 5.7 mV) at pH=7.4, was due to the presence of positively charged-NH3^+^ end groups of the grafted PEG (Table 1). The NF@PEG and NF@SiO_2_ @PEG exhibited respectively a saturation magnetization (Ms) of 356 and 340 kA/m respectively, close to the Ms value of bulk maghemite γ-Fe_2_O_3_ (~400 kA/m) (Figure 1C-D, Table 1). The difference between the Ms of both type of nanoparticles was very low, indicating that the SiO_2_ shell did not modify the saturation magnetization of NF@SiO_2_@PEG comparatively to NF@PEG. Moreover, the absence of hysteresis in the M vs *H* curves when the field intensity decreases indicated that NF@PEG and NF@SiO_2_@PEG exhibited a superparamagnetic behavior at 293K. The M vs *H* curves fitting with the Langevin function convolved with a log-normal distribution of nanoparticles core diameters and showed very similar mean number-averaged (d^n^) and volume-averaged (d^w^) magnetic size (Figure 1C-D, Table 1). The heat efficiency of NF@PEG and NF@SiO_2_@PEG, measured through the AC magnetometry, showed that NF@SiO_2_@PEG presented a lower specific absorption rate (SAR) determined from area of the dynamic hysteresis loop, than NF@PEG (Figures 1E and S3). The SAR of NF@PEG and NF@SiO_2_ @PEG, determined at the conditions close to the ones used *ex vivo* (19 kA.m^−1^, 275 kHz), was respectively 186.0 ± 5.9 and 180.0 ± 4.9 W/g γ-Fe_2_O_3_ (Figure 1E). Calorimetric assay showed similar SAR for both types of nanoparticles, measured at 93 kHz as a function of the magnetic field amplitude (Figure 1F). All together, these results showed that the surface state, the magnetic properties and the heating efficiency under AMF of both NF@PEG and NF@SiO_2_@PEG were very comparable.

**Table 1.**

|  | NF@PEG | NF@PEG@Gastrin | NF@SiO2@PEG | NF@SiO2@PEG@Gastrin |
| --- | --- | --- | --- | --- |
| DTEM (nm) | 17.3 ± 2.6 |  | 23.4 ± 3.7<br>(SiO2 shell 3.1 ± 0.2) |  |
| DH (nm) | 50.3 ± 4.0 | 75.8 ± 5.1 | 64.5 ± 6.9 | 79,3 ± 7.3 |
| PDI | 0.25 ± 0.15 | 0.15 ± 0.01 | 0.17 ± 0.02 | 0.15 ± 0.03 |
| ζ (mV) | +14.0 ± 2.0 | -21,6 ± 1.1 | +22.6 ± 5.7 | -1.0 ± 0.5 |
| RF/D (PEG regimen) | 2.4 (brush) |  | 2.7 (brush) |  |
| Ms (kA.m<sup>-1</sup>) | 356 |  | 340 |  |
| Dn (nm) | 9.1 ± 0,8 |  | 10.2 ± 0,9 |  |
| Dw (nm) | 11.2 ± 0,7 |  | 11.9 ± 1,1 |  |

**Figure 1.**
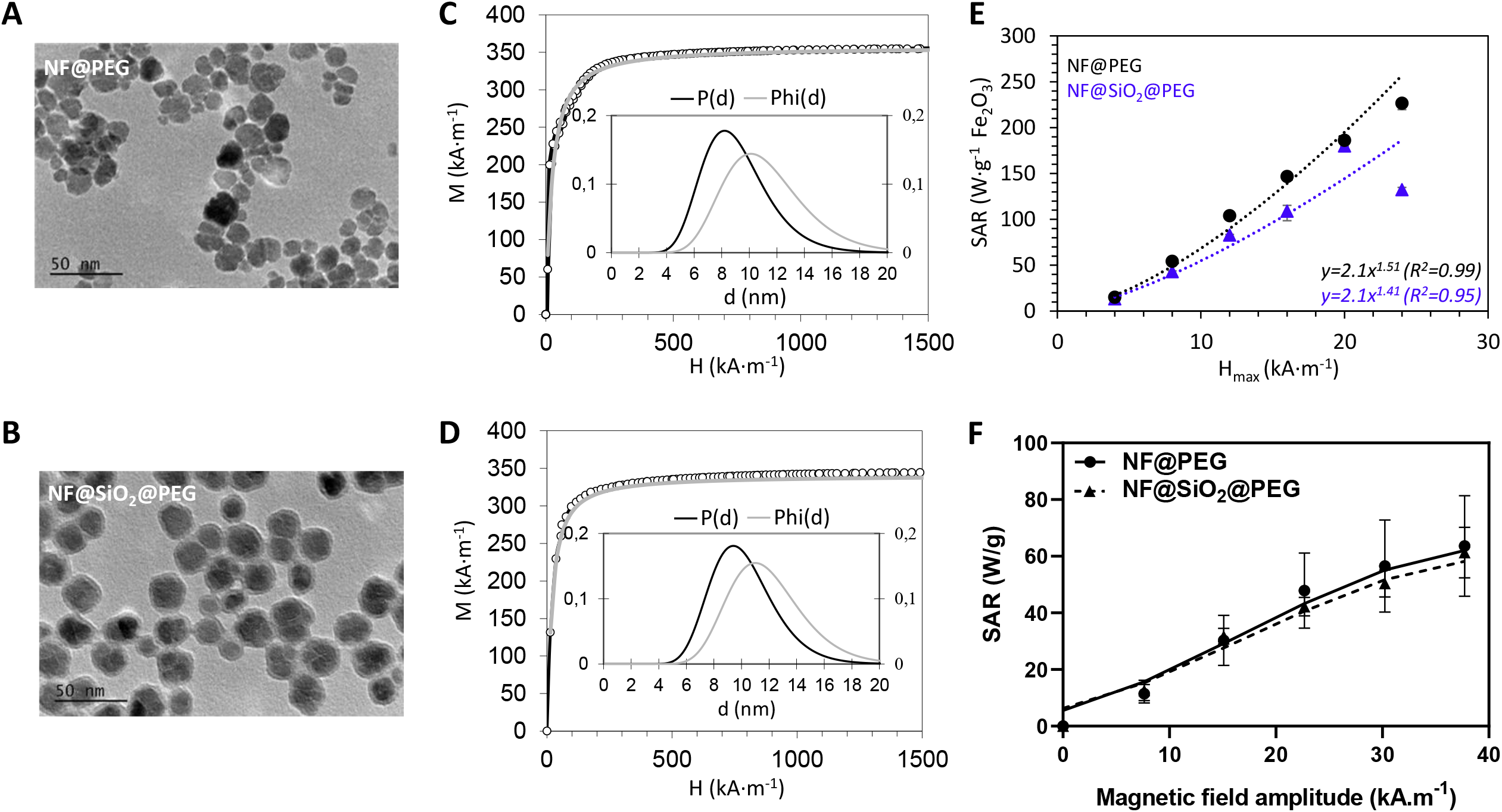
Physico-chemical properties of NF and NF@SiO_2_ iron oxide MNPs. **(A,B)** TEM images of NF@PEG (A) and NF@SiO_2_@PEG (B), **(C,D)** M vs H curves under DC field at 300 K of NF@PEG (C) and NF@SiO_2_@PEG (D) both fitted by Langevin’s law of superparamagnetism, corresponding to the grey lines. The curves in inset correspond to the magnetic size distribution laws obtained from the Langevin fit of magnetization convolved with a distribution law *Phi*(*d*) of the volume fraction of the nanoparticles associated to a log-normal law *P*__(*d*_0,*σ*) (*d*) for their number distribution. **(E)** Specific absorption rate (SAR) of a NF@PEG and NF@SiO_2_@PEG solution (3 mg γ-Fe_2_O_3_ /ml) measured at a frequency of 275 kHz as a function of the magnetic field amplitude, through AC magnetometry. Error bars correspond to three separate experiments. **(F)** Specific absorption rate of a NF@PEG and NF@SiO_2_@PEG solution (6 mg γ-Fe_2_O_3_ /ml) measured at a frequency 93 kHz as a function of the magnetic field amplitude, through calorimetric assay. Results are expressed as mean ± sem of three separate experiments

The degradation of NF@PEG and NF@SiO_2_@PEG and the temperature variations A were then analyzed in a pH4.5 medium mimicking the lysosomal environment, named as Artificial Lysosomal Fluid (ALF) ^29^, upon AMF exposure (21 kA m^−1^, 217 kHz) or not (Figures 2A and S4). Different concentrations of NF@PEG and NF@SiO_2_@PEG in ALF were thermalized at 37 °C, then exposed or not to AMF during 2h. Fe^3+^ release from γ-Fe O dissolution was analyzed in the supernatant by UV-VIS spectroscopy. The results showed that AMF exposure increased Fe^3+^ release from NF@PEG but not from NF@SIO_2_@PEG whatever the nanoparticles concentration, indicating that the silica shell acts as a protective physical diffusion barrier that prevented the degradation of γ-Fe_2_0_3_ core and Fe^3+^ release during the experimental range (Figures 2A and S4). Once the magnetic field was applied, the temperature of the NF@PEG and NF@SiO_2_@PEG colloidal solutions rapidly increased from 37 °C to 45, 65 or 92 °C for NF@PEG and NF@SiO_2_@PEG at 0.5, 3 and 7 g/L γ-Fe_2_O_3_ respectively, as a result of magnetic hyperthermia (Figure S4A). At 0.5 and 3 g/L γ-Fe_2_O_3_, the temperature of NF@PEG and NF@SiO_2_@PEG solutions remained constant throughout all the duration of AMF exposure, 45 and 65°C respectively, and a progressive degradation of NF@PEG (10.7 ± 1.0 % and 26.6 ± 1.3 %, respectively) was observed whereas the degradation of NF@SiO_2_@PEG was negligible (1.6 ± 0.9 % and 0.5 ± 0.1 %, respectively) (Figures 2A and S4B-C). Of note, water bath incubation (at the temperature corresponding to AMF application on the IONP solutions) for 2h induced similar Fe release from NF@PEG (11.2 ± 0.9 % vs 13.4 ± 0.8 % at 0.5 g/L γ-Fe_2_O_3_; 26.6 ± 1.3 % vs 20.4 ± 0.6 at 3 g/L, Figure S4B-C) than AMF exposure, showing that the NF@PEG dissolution upon AMF application was a result of macroscopic temperature increase in the tube. At 7 g/L γ-Fe_2_O_3_, AMF exposure induced a high degradation of NF@PEG with 79.1 ± 0.4 % of Fe^3+^ released in the supernatant (2.2 ± 0.1 % in absence of AMF), whereas NF@SiO_2_@PEG remained intact (1.4 ± 0.2 % and 0.5 ± 0.1 % Fe release in presence or absence of AMF respectively, Figure 2A). Interestingly, the temperature of the NF@PEG and NF@SiO_2_@PEG solutions at 7 g/L γ-Fe_2_O_3_ reached 92°C under AMF application. However, the temperature of the NF@SiO_2_@PEG solution remained constant throughout all the duration of AMF exposure, whereas the one of NF@PEG solution decreased after around 20 minutes of AMF application to a second temperature plateau at 60°C, suggesting that the decrease in the heating power of NF@PEG leading to the decrease of macroscopic temperature of the NF@PEG solution during AMF application could be attributed to the gradual dissolution of the iron oxide core (Figure S4A). This degradation of the NF@PEG in ALF induced by AMF exposure was also confirmed by TEM analysis (Figure S5). The diameter of the NF@PEG magnetic core decrease from 17.3 ± 2.6 nm to 12.4 ± 2.5 nm and 10.7 ± 2.4 nm after 1h and 2h under AMF. As expected, the diameter of the NF@SiO_2_@PEG magnetic chore was not affected by AMF application. All together, these results indicated that NF@PEG have degraded and released Fe^3+^ ions in acidic medium upon AMF exposure, and that degradation rate of NF@PEG increased with the nanoparticles concentration. The kinetics of Fe^3+^ release from NF@PEG induced by AMF also indicated that NF@PEG degradation was more rapid and important with highest NF@PEG concentration (Figures S4D-F). In contrast, the degradation of NF@SiO_2_@PEG core was not measurable for the different tested concentrations, demonstrating the effectiveness of the silica shell as protective barrier against γ-Fe_2_O_3_ magnetic core dissolution in acidic medium under AMF. These results are in agreement with a previous study showing that an uninterrupted gold shell could preserve the IONPs magnetic core and properties, confirming the feasibility of this strategy ^18^.

**Figure 2:**
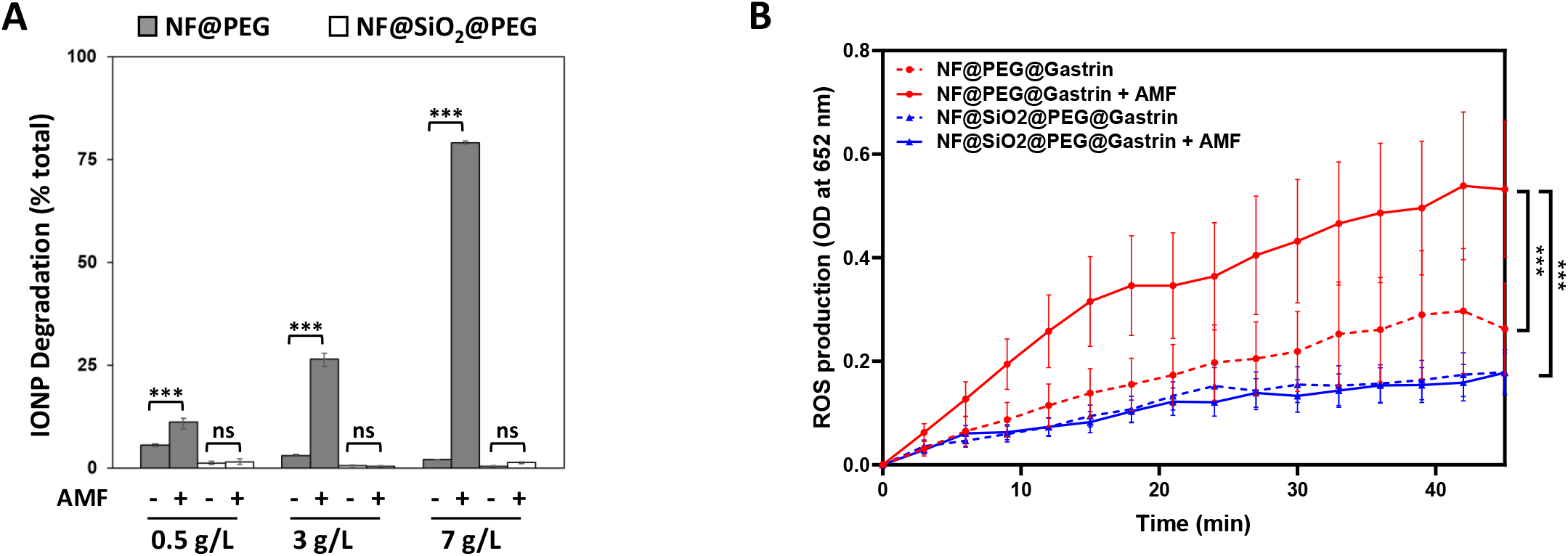
**(A)** *In vitro* Fe dissolution from IONPs. NF@PEG and NF@SiO_2_@PEG solutions in Artificial Lysosomal Fluid (ALF) were exposed or not to AMF during 2h, then centrifuged (60,000g, 25 min). Fe dissolution was analyzed in the surpernatant by UV-VIS spectroscopy. **(B)** *In vitro* ROS production by NF@PEG and NF@SiO_2_@PEG exposed or not AMF. 10 µg NF@PEG and NF@SiO_2_@PEG were incubated 72h in pH 4.4 medium, then exposed or not to AMF for 2h with temperature maintained at 37°C. H_2_O_2_ and TMB were then added and optical absorbance was analyzed by spectrometry at 652 nm as a function of time.

### Silica shell prevents *in vitro* catalysis of ROS by magnetic hyperthermia

Fe cation and iron oxide nanoparticles were shown to present peroxidase-like activity, which catalyze the hydrogen peroxide (H_2_ O_2_) to generate highly cytotoxic hydroxyl radical (^•^OH) through Fenton and Haber-Weiss reactions ^30,31^. In order to evaluate the efficacy of NF@PEG and NF@SiO_2_ @PEG in producing hydroxyl radicals, we employed the colorimetric method based on the oxidation of 3,31.,5,51.-tetramethylbenzidine (TMB) in the presence of H_2_O_2_ under acidic conditions mimicking lysosome environment. In the presence of H O and AMF application (21 kA.m^−1^, 275 kHz), NF@PEG increased the oxidation of TMB to form the oxidized and therefore blue-colored TMB (oxTMB), measured at the characteristic absorbance 652 nm^30,31^, comparatively to NF@PEG in absence of AMF application, indicating that unprotected NF@PEG enhanced the conversion of H_2_ O_2_ to ^•^OH during magnetic hyperthermia (Figure 2B). In contrast, negligible absorbance was observed in the presence of NF@SiO_2_@PEG, suggesting that NF@SiO_2_@PEG do not promote the oxidation of TMB, in response or not to AMF application (Figure 2B). This result demonstrates that the SiO_2_ layer was an efficient shield that protected efficiently the γ-Fe_2_O_3_ magnetic core preventing Fe cations release from the nanoparticles and the production of hydroxyl radicals from H_2_O_2_. Moreover, these results indicate that NF@PEG peroxidase-like activity may involve the Fe cations released from the degradation of the unprotected γ-Fe_2_O_3_ magnetic core or be catalyzed at the surface of the unprotected magnetic core. Thus, hydrated PEG corona does not constitute a barrier preventing the Fe^3+^ release from γ-Fe_2_O3 magnetic core and/or the catalysis of peroxidase-like activity at the γ-Fe2O3 magnetic core surface of NF@PEG.

### Nanoparticles uptake, subcellular localization and intracellular heating

To investigate the behavior of NF@PEG and NF NF@SiO2@PEG to generate intracellular ^•^OH and induce cell mortality under magnetic hyperthermia experiments, we first conjugate both NF@PEG and NF@SiO2@PEG with a synthetic peptide analog of gastrin according a procedure previously described ^32^, to increase their uptake by pancreatic cancer cells expressing the type 2 cholecystokinin receptor (CCK2R), chosen as a model, and their accumulation in the lysosomes. NF@PEG and NF@SiO2@PEG were also decorated with a fluorophore (DY647), allowing their fluorescence detection ^32^. The hydrodynamic diameter of NF@PEG@Gastrin and NF@SiO2@PEG@gastrin, measured in PBS pH7.4, were respectively of 75.8 ± 5.1 and 79.3 ± 7.3 nm, while their zeta potential values measured in 1 mM HEPES at pH 7.4 were −21.6 ± 1.1 and −1.0 ± 0.5 mV (Table 1). We then analyzed the cytotoxicity, the uptake and subcellular localization of these nanoparticles in MiaPaca-2 cells expressing or not the CCK2 receptor. NF@PEG@Gastrin and NF@SiO2@PEG@Gastrin do not present cytotoxic activity on MiaPaca-2 and MiaPaca2-CCK2 cells up to 128 µg γ-Fe_2_O_3_/ml (Figure S6). The hemocompatibility was also investigated, and the hemolysis ratios of the red blood cells for NF@PEG and NF@SiO2@PEG were negligible at concentrations up to 345 µg γ-Fe_2_O_3_, indicating that these magnetic nanoparticles are biocompatible for intravenous administration (Figure S7).

The kinetics of uptake of both modified IONPs (NF@PEG@Gastrin and NF@SiO2@PEG@Gastrin) determined by flow cytometry indicate that their amount taken up by MiaPaca2-CCK2 increased with the time of incubation (Figures 3A-B). Furthermore, both IONPs bind and internalize much higher and faster in MiaPaca2-CCK2 than in MiaPaca2 cells, indicating that gastrin improves the uptake kinetics of both nanoparticles type in MiaPaca2-CCK2 cells thanks to specific and high affinity binding of gastrin to the CCK2 receptor. Indeed, the uptake of NF@PEG@Gastrin and NF@SiO2@PEG@Gastrin is respectively 3.7 ± 1.0 and 2.0 ± 0.2-fold more important in MiaPaca2-CCK2 cells compared to MiaPaca2 cells, after 72 hours of incubation. The targeting efficiency of NF@Gastrin was also determined by flow cytometry after 72h of incubation and confirmed that NF@PEG@Gastrin and NF@SiO2@PEG@Gastrin uptake is more important in cells expressing the CCK2 receptor than control cells, at any nanoparticles concentrations, confirming that the use of gastrin is efficient to target pancreatic cancer cells expressing the CCK2 receptor (Figure S8). The absolute amount NF@PEG@Gastrin and NF@SiO2@PEG@Gastrin internalized by MiaPaca2-CCK2 cells was then determined by NMR relaxometry, being in the same order for both nanoparticles: 2.8 ± 0.7 (T1) / 2.9 ± 0.7 (T2) pg γ-Fe_2_O_3_ / cell and 3.0 ± 1.6 (T1) / 3.1 ± 2.0 (T2) pg γ-Fe_2_O_3_ / cell incubated for 72h with NF@PEG@Gastrin or NF@SiO2@PEG@Gastrin, respectively (Figure 3C). Then, we characterized the cellular fate of NF@PEG@Gastrin or NF@SiO2@PEG@Gastrin in MiaPaca2-CCK2 (Figure 4). The transmission electron microscopy (TEM) images of MiaPaca2-CCK2 cells incubated with both IONPs were mainly localized in the lysosomes of MiaPaca2-CCK2, after 72h of incubation (figures 4 A and B). Confocal microscopy study confirmed that both IONPs accumulated in the lysosomal vesicles (Figures 4C-D). Analysis of IONPs colocalization with Lysotraker demonstrated that 78.4 ± 1.2 and 78.4 ± 1.0 % of internalized NF@PEG@Gastrin or NF@SiO2@PEG@Gastrin were localized inside the lysosomes of MiaPaca2-CCK2 cells, respectively (Figures 4E).

**Figure 3.**
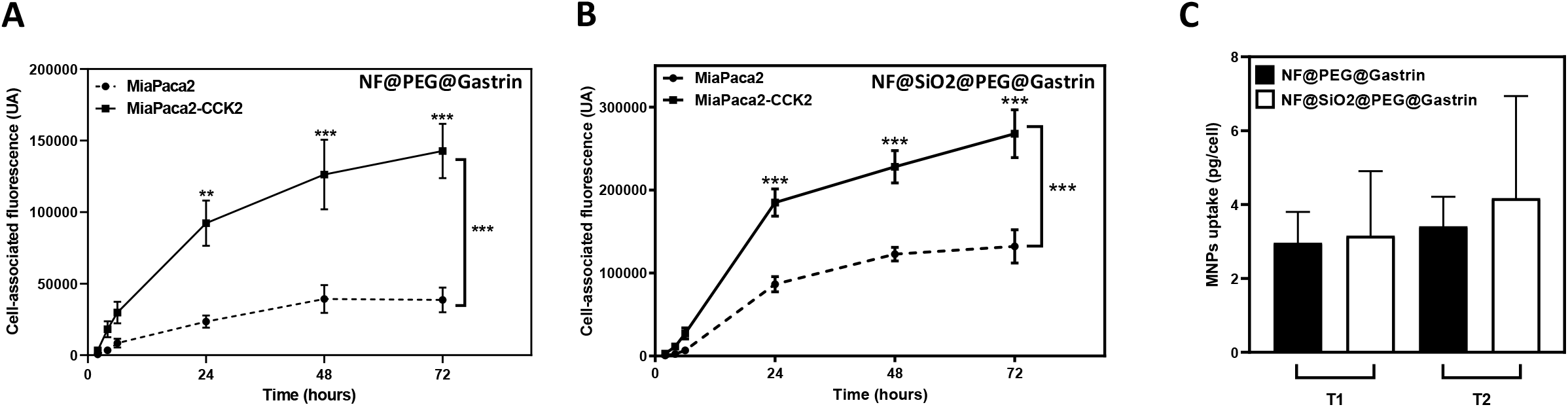
Analysis of nanoparticles uptake. Kinetics of NF@PEG@Gastrin **(A)** and NF@SiO2@PEG@Gastrin **(B)** uptake. MiaPaca2 and MiaPaca2-CCK2 cells were incubated with 16 µg/ml of fluorescent NF@PEG@Gastrin or NF@SiO2@PEG@Gastrin for 2, 4, 6, 24, 48 or 72h at 37°C. Fluorescence was measured by flow cytometry, and results are expressed as fluorescence associated with the cells and are the mean ± SEM of at least four separate experiments. **(C)** Analysis of MNPs uptake by relaxometry. MiaPaca2-CCK2 cells were incubated with 16 µg/ml of NF@SiO2@PEG@Gastrin for 72h at 37°C, centrifuged.

**Figure 4.**
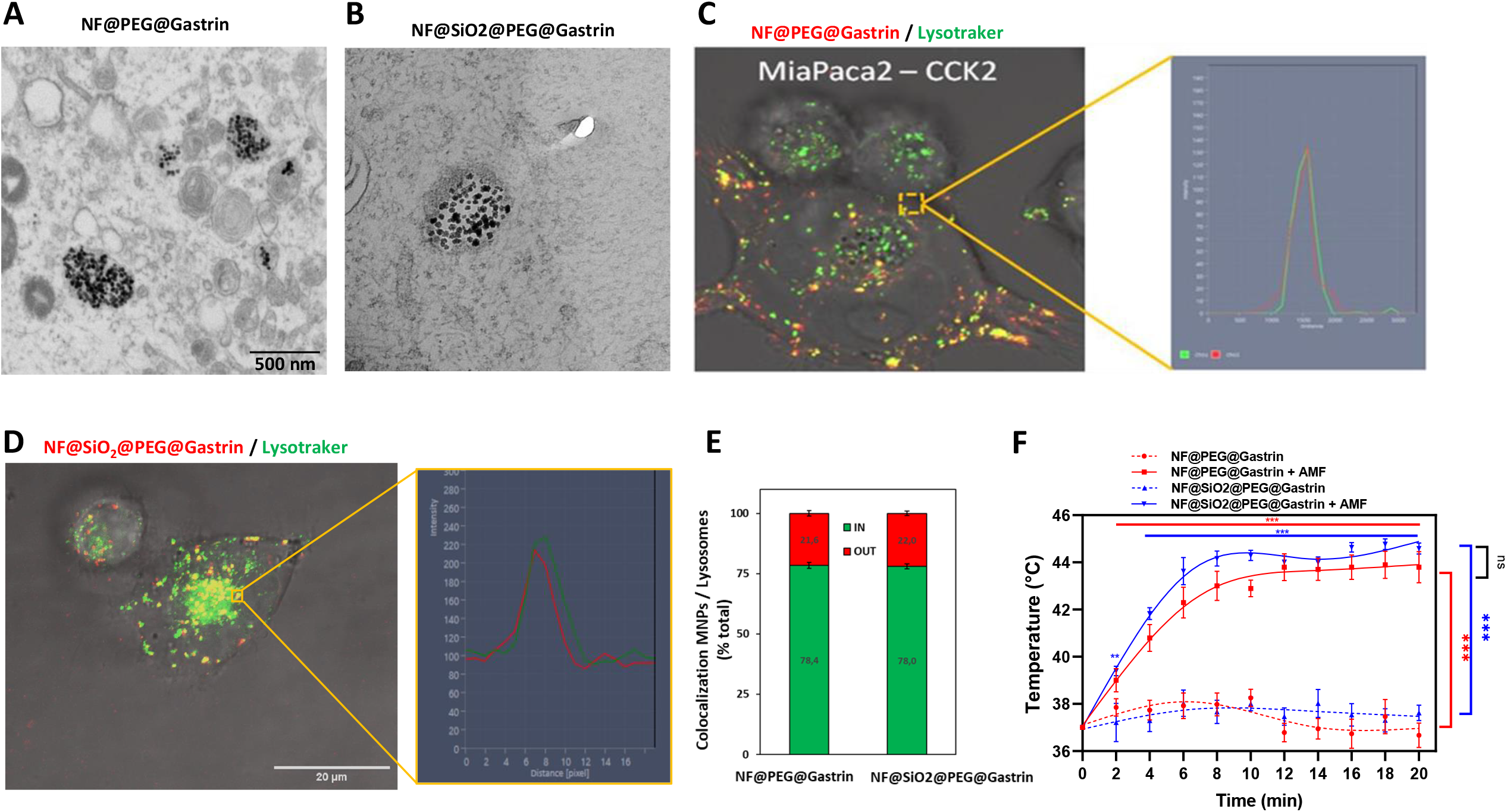
Subcellular localization of MNPs. Electron microscopy analysis of NF@PEG@Gastrin **(A)** and NF@SiO2@PEG@Gastrin **(B)** accumulation in MiaPaca2-CCK2 cells. Image representative of MNPs accumulated in in lysosomes, (A) and (B). Analysis of NF@PEG@Gastrin **(C)** and NF@SiO2@PEG@Gastrin **(D)** localization by confocal microscopy. MiaPaca2-CCK2 cells were incubated with 16 µg/ml of MNPs for 72 h, then incubated with 10 nM Green Lysotraker and observed by confocal microscopy. **(E)** Quantification of MNPs accumulation in lysosomes by analysis of confocal microscopy images: green = inside lysosomes, red = outside lysosomes. MiaPaca2-CCK2 cells were incubated with 16 µg/ml of MNPs for 72 h. **(F)** MiaPaca2-CCK2 cells with NF@PEG@Gastrin or NF@SiO2@PEG@Gastrin decorated with DY549. AMF was applied using a miniaturized electromagnet. Temperature at MNPs surface was analyzed by measuring the decrease of theDY549 fluorescence intensity and calculated using the calibration curve (Figure S9). 20–30 cells/experiments were analyzed and results are the mean ± SEM. Significant differences comparatively to t = 0 are indicated above the curve; significant differences between conditions are indicated with brackets.

We also evaluated the temperature rise at the surface of both nanoparticles during MILH experiments, using the fluorescent molecular thermometers DY549 attached on the PEG corona as previously described ^23^. Quantitative fluorescence imaging was then carried out on cells exposed or not to AMF for a limited duration (⍰20-min) and incubated with ROS scavenger NAC in order to preserve lysosome integrity and avoid possible chemical quenching of the fluorescent molecular thermometers by ROS. We showed that the DY549 fluorescence intensity declined similarly for both nanoparticles with the time of AMF exposure, while it did not vary in absence of AMF (Figure S9B). According to the calibration curve (Figure S9A), the percentage of decrease of fluorescence intensity observed after ~6-min of AMF application corresponds respectively to a temperature increase of 6.0 ± 0.9 and 6.6 ± 0.6 °C at the surface of NF@PEG@Gastrin or NF@SiO2@PEG@Gastrin, indicating that the temperature actually reached ~43 at their surface (Figure 4F).

Taken together, these results indicate that both nanoparticles had similar *ex vivo* behavior since NF@PEG@Gastrin and NF@SiO_2_@PEG@Gastrin as they preferentially internalized in MiaPaca2-CCK2 cells and accumulated in their lysosomes. Moreover, both nanoparticles present similar heating capacity in cells, allowing the comparison of their performance during magnetic hyperthermia experiments.

### Analysis of NF@PEG@Gastrin and NF@SiO2@PEG@Gastrin efficacy as anticancer agent by magnetic hyperthermia

Although we previously showed that cell death depends on ROS production by Fenton reaction, the origin of Fe cations, critical to produce ROS in MILH, is still elusive ^11^. In the literature, Fe cations were assumed to be released by IONPs surface resulting in a greater extent of ROS catalysis in acidic medium such as lysosome compartment (ref). To gain insight into the cell death mechanism, we thus investigated the efficacy of NF@PEG@Gastrin and NF@SiO2@PEG@Gastrin as anticancer agent by MILH. The effect of AMF application to induce the death of MiaPaca2-CCK2 cells having internalized NF@PEG@Gastrin or NF@SiO2@PEG@Gastrin was first evaluated. For this purpose, MiaPaca2-CCK2 cells were incubated with increasing concentration of NF@PEG@Gastrin or NF@SiO2@PEG@Gastrin for 72h, washed to eliminate unbound and non-internalized nanoparticles and exposed to AMF (21 kA.m^−1^, 275 kHz) for 2h. Surprisingly, both IONPs induce cell mortality with the same efficacy, by MILH so without any perceptible temperature increase (Figure 5A). The effect of MILH on cell viability was also studied by exposing cells for 2h three times (days 5, 7 and 9) and showed that both NF@PEG@Gastrin and NF@SiO_2_@PEG@Gastrin inhibit the cell proliferation with the same efficacy (Figure 5B). Cell mortality induced by NF@SiO_2_@PEG@Gastrin may at first suggest that SiO_2_ coating does not ensure the complete protection of IONP core in cells, and that it may be degraded in the endo-lysosomal compartment releasing of Fe^3+^ ions similarly as for NF@PEG@Gastrin. However, we previously demonstrated that SiO_2_ shell protects the iron oxide core and prevents the release of Fe cations even under highly dissociating conditions such as AMF application or water bath increasing the temperature up to 92°C, in ALF acidic medium (Figures 2 and S4). Moreover, the kinetics of IONPs degradation was shown to be slowed down in the endo-lysosomal compartment of cultured cancer cells or murine liver and spleen, comparatively to *in vitro* lysosome-like medium, due to IONPs aggregation that may protect them from the degradation ^16^. The alternative explanation is that protected NF@SiO_2_@PEG@Gastrin induce cell mortality and inhibit cell viability by the local release of thermal energy under AMF without involving Fe^3+^ liberation from IONPs degradation. We thus investigated whether NF@SiO_2_@PEG@Gastrin enhanced the production of ROS upon AMF application. As shown in Figure 5C, the protected NF@SiO_2_@PEG@Gastrin increased the production of ROS with the same efficacy than NF@PEG@Gastrin under MILH (increased ROS levels by 4.4 ± 0.5 and 3.4 ± 0.5-fold compared to control cells, respectively). Moreover, this effect was associated with an increased permeabilization of lysosomes (Figure 5D). All together, these results demonstrated that both unprotected NF@PEG@Gastrin and protected NF@SiO_2_@PEG@Gastrin IONPs induced similarly the production of ROS leading to lysosome membrane permeabilization and cell death. It should be highlighted that lysosomes contain endogenous iron, mainly in the form of Fe^2+^, as a result of degradation of iron-containing proteins ^33,34^. Thus, in absence of Fe liberation from protected NF@SiO_2_@PEG@Gastrin IONPs, local heat release by NF@SiO_2_@PEG@Gastrin IONPs during MILH enhances the ROS production through Fenton reaction using endogenous Fe^2+^. Concerning the NF@PEG@Gastrin, we cannot exclude that these unprotected IONPs increases ROS production upon MILH through two parallel routes: the local heat release from IONPs that induce ROS production through Fenton reaction using endogenous Fe^2+^ as well as from dissolution of Fe^3+^ ions from NF degradation.

**Figure 5.**
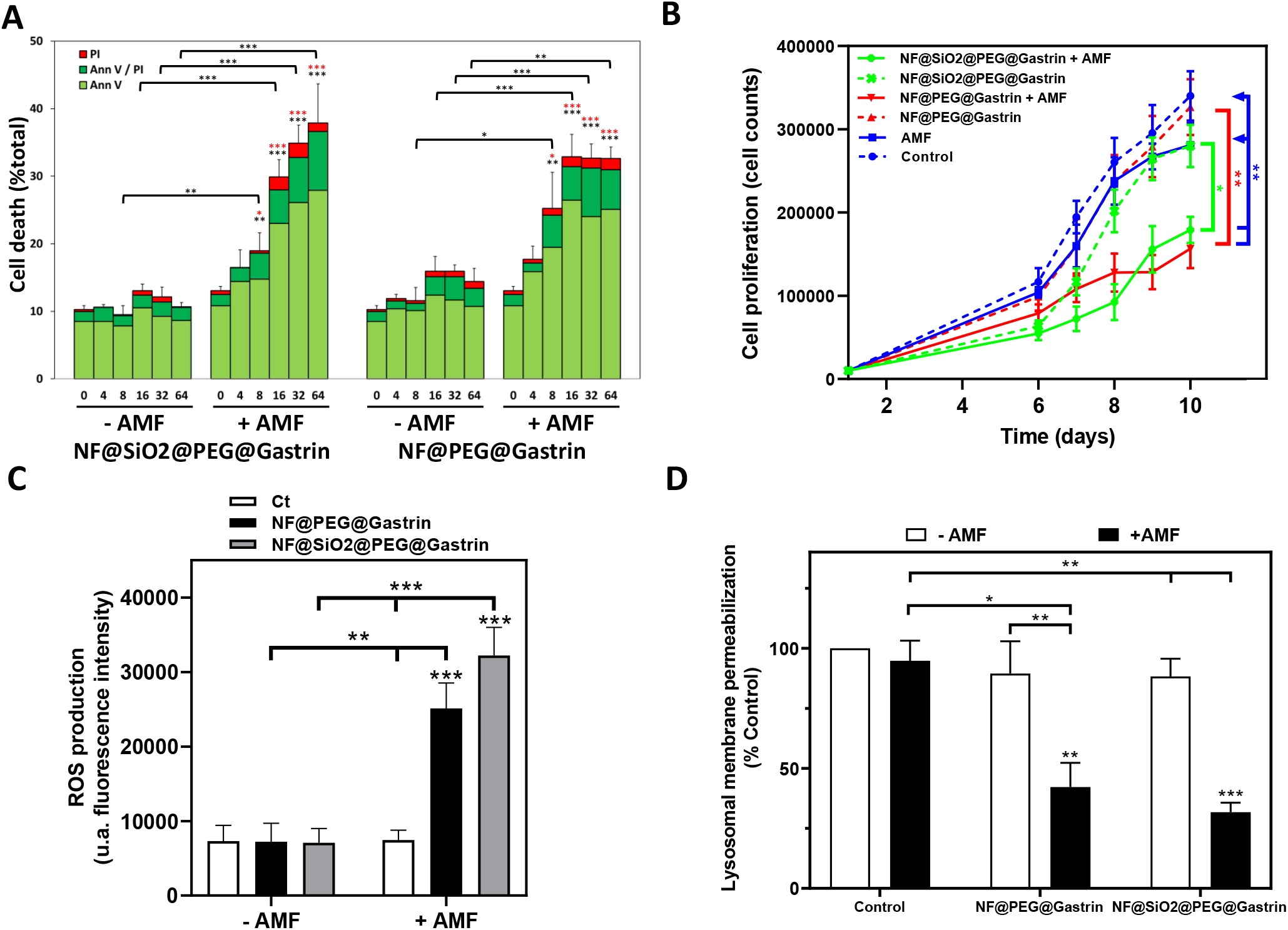
Bilogical impacts of Magnetic hyperthermia induced by NF@PEG@Gastrin or NF@SiO2@PEG@Gastrin. Cells were incubated or not with 16 µg/ml NF@PEG@Gastrin or NF@SiO2@PEG@Gastrin for 72h, then exposed or not to AMF for 2h. **(A)** ROS production was analyzed using CellROX Green, immediately after AMF exposure, by analyzing the fluorescence intensity from confocal microscopy images and expressed as fold change of fluorescence intensity overcontrol cells (in absence of nanoparticles and AMF). 2000–3000 cells/experiments were analyzed, and results are the mean ± SEM of 4 separate experiments. Statistical analysis was performed using two-way ANOVA test: significant difference compared to the control condition (cells devoid of nanoparticles in absence of AMF exposure) indicated above histogram bar, statistical significances between other conditions indicated by lines/brackets. **(B)** After AMF application, cells were incubated with 10 nM Green Lysotraker and observed by confocal microscopy. Lysosomal membrane permeabilization was analyzed by measuring the Lysotraker fluorescence intensity from confocal microscopy images and expressed as a % of fluorescence intensity overcontrol cells (in absence of nanoparticles and AMF). 2000–3000 cells/experiments were analyzed, and results are the mean ± SEM of 4 separate experiments. Statistical analysis was performed using two-way ANOVA test: significant difference compared to the control condition (cells devoid of nanoparticles in absence of AMF exposure) indicated above histogram bar, statistical significances between other conditions indicated by lines/brackets. **(C)** Dead cells were labeled with AnnV/PI and counted 4 h after AMF exposure by confocal microscopy. Results are the mean ± SEM of 4 separate experiments. Statistical analysis was performed using one-way ANOVA test: significant difference compared to the control condition (cells devoid of nanoparticles in absence of AMF exposure) or AMF condition (without nanoparticles) indicated respectively in black or red above histogram bar, statistical significances between other conditions indicated by lines/brackets. **(D)** Cells incubated or not with nanoparticles were exposed to AMF three times (days 5, 7 and 9) for 2 h. Cell proliferation was evaluated by counting cells each day. Results are expressed as mean ± SEM of at least four separate experiments. Statistical analysis was performed using one-way ANOVA test.

### Impact of endogenous Fe depletion on ROS production and cell death induced by magnetic hyperthermia

To elucidate the role of endogenous Fe^2+^ in the cell death mechanism, MILH experiments were conducted in the presence of Ferristatin-II, which inhibits iron uptake by the cell by promoting the degradation of transferrin receptor 1 (TfR1) ^25,35^. Iron transportation in cells involves transferrin (Tf) and TfR1. Iron bound to Tf/TfR1 is transported to cells and internalizes through receptor-mediated endocytosis. The iron is then released from Tf in the acidic endo-lysosomal compartment. Following the aforementioned studies, MiaPaca2-CCK2 cells were treated with 50µM Ferristatin-II for 24h in SVF-free medium. Fe depletion in the cells was then evidenced by confocal microscopy using FerroOrange, a dye that detects and shows strong fluorescence signal upon reaction with free Fe^2+ 36^. As expected, the quantity of Fe^2+^ is significantly and strongly reduced in the cells treated with 50 µm Ferristatin-II for 24h, compared to control cells in absence of Ferristatin-II, indicating that Ferristatin-II depleted the Fe^2+^ content in the MiaPaca2-CCK2 cells (Figure 6A). MILH experiments were then conducted on NF@PEG@Gastrin or NF@SiO2@PEG@Gastrin-loaded cells in the presence or not of Ferristatin-II. For this, MiaPaca2-CCK2 cells were incubated with NF@PEG@Gastrin or NF@SiO2@PEG@Gastrin for 72h allowing their accumulation in the lysosomes, washed and incubated with 50 µM Ferristatin-II in SVF-free medium for 24h. After AMF application, ROS production was then analyzed. We showed that Ferristatin-II prevent MILH-increased ROS production which returned to the basal value in NF@SiO2@PEG@Gastrin-loaded cells, indicating Fenton reaction producing ROS can be carried out using endogenous Fe^2+^ and that local heat release from IONPs upon AMF application promote ROS overproduction leading to cell death (Figure 6B). Surprinsingly, ROS production is also prevented by Ferristain-II in NF@PEG@Gastrin-loaded cells during MILH, although these unprotected IONP are prone to degradation, strongly suggesting that Fe cations provided from NF@PEG@Gastrin degradation induced by AMF do not constitute the main source of Fe ion to generate ROS during MILH (Figure 6C). Moreover, these results highlighted that the endogenous Fe^2+^ is crucial to enhance the production of ROS during MILH. Although the unprotected NF@PEG@Gastrin surely released Fe^3+^ ions during their trafficking in the endo-lysosomal compartment and MILH, the local heat generated by MILH is sufficient to generate ROS by upregulating the Fenton reaction involving the endogenous Fe^2+^ ions. We further analyzed the impact of Ferristatin-II treatment on MILH efficacy to induce cell death in NF@PEG@Gastrin or NF@SiO2@PEG@Gastrin-loaded cells (Figure 7). We found that the addition of Ferristatin-II, which deplete cells in iron and prevent ROS production during MILH, inhibits cell death induced by MILH in both in NF@PEG@Gastrin and NF@SiO2@PEG@Gastrin-loaded cells. As a control, we showed that Ferristatin-II does not inhibit the gemcitabine-induced cell death (Figure S10). All together, these results demonstrate that MILH engender ROS production through Fenton reaction using endogenous Fe^2+^ rather than the Fe^3+^ released from IONPs degradation, leading to cancer cell death.

**Figure 6.**
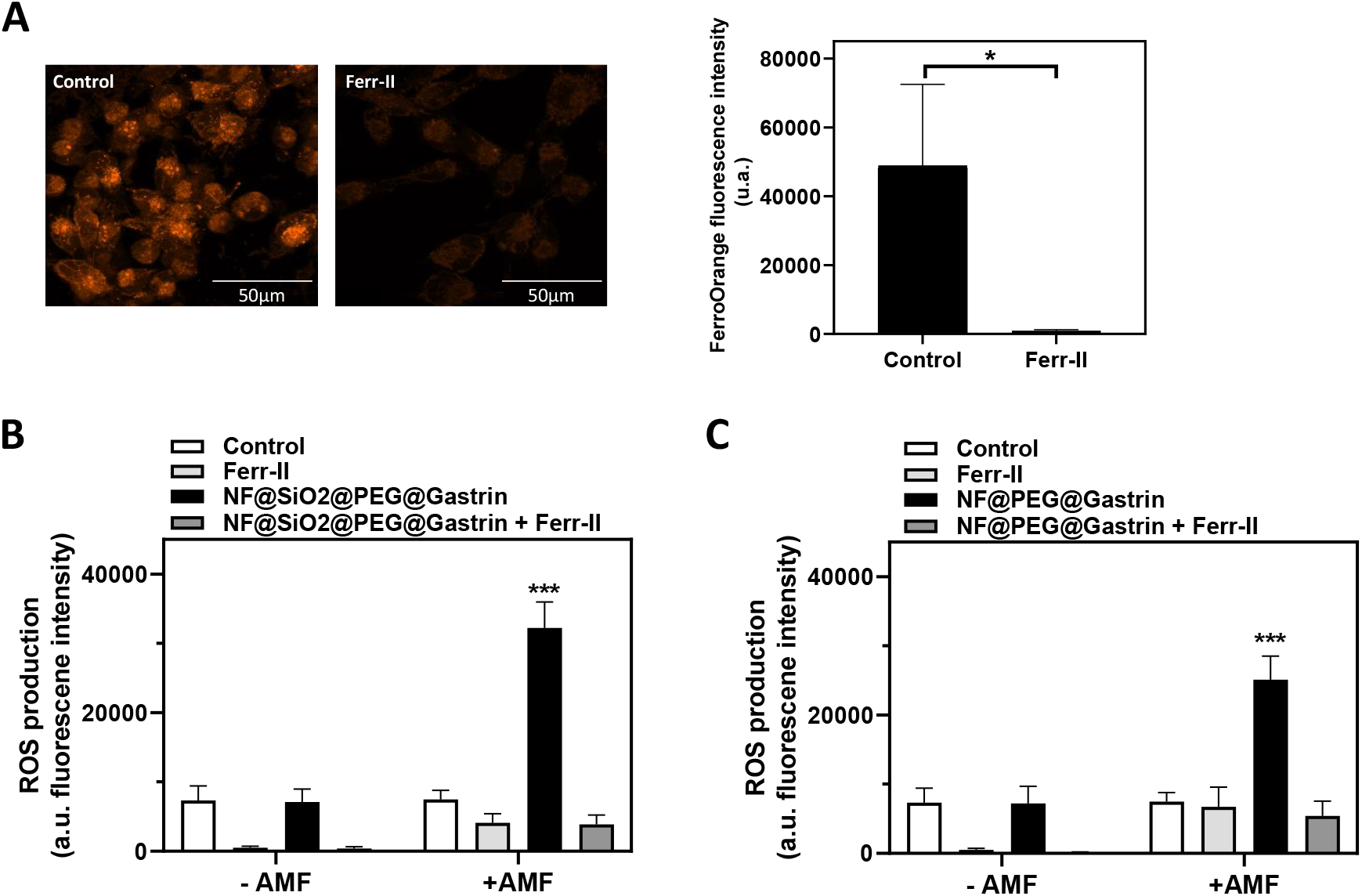
Impact of endogenous Fe depletion on ROS production induced by magnetic hyperthermia. **(A)** Endogenous Fe depletion by Ferristatin-II treatment. Cells were incubated with 50µM Ferristatin-II (Ferr-II) in DMEM medium without SVF for 24h, then stained with FerroOrange. FerroOrange fluorescence intensity was analyzed from confocal microscopy images (2000–3000 cells/experiments). Results are the mean ± SEM of 4 separate experiments. Statistical analysis was performed using t-test. **(B, C)** Cells were incubated or not with 16 µg/ml NF@PEG@Gastrin or NF@SiO2@PEG@Gastrin for 72h, then incubated or not with Ferr-II in DMEM medium without SVF for 24h and exposed or not to AMF for 2h. ROS production was analyzed using CellROX Green, immediately after AMF exposure, by analyzing the fluorescence intensity from confocal microscopy images. 2000–3000 cells/experiments were analyzed, and results are the mean ± SEM of 4 separate experiments. Statistical analysis was performed using two-way ANOVA test: significant difference compared to all other conditions indicated above histogram bar.

**Figure 7.**
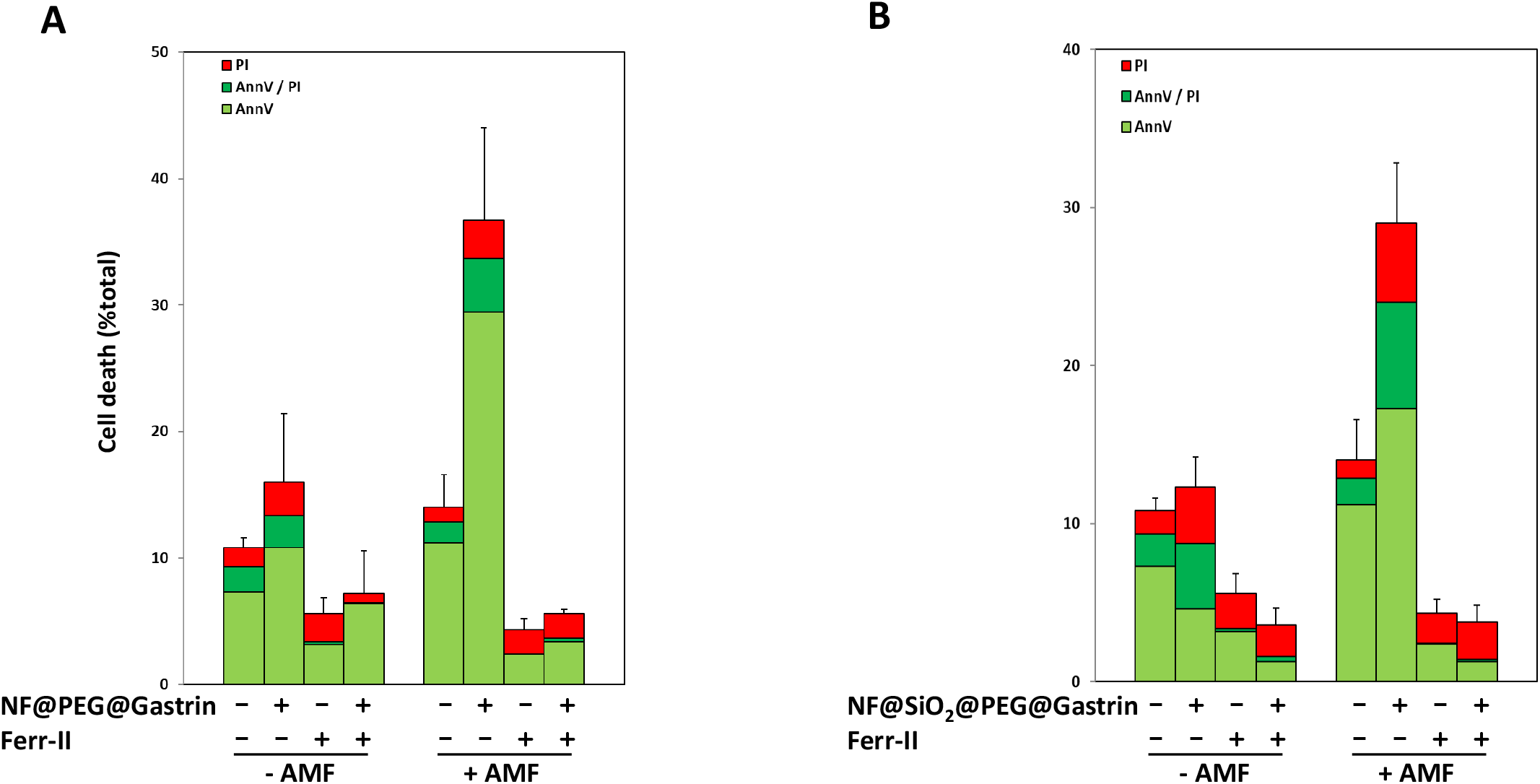
Impact of endogenous Fe depletion on cell death induced by magnetic hyperthermia. Cells were incubated or not with 16 µg/ml NF@PEG@Gastrin or NF@SiO2@PEG@Gastrin for 72h, then incubated or not with Ferr-II in DMEM medium without SVF for 24h and exposed or not to AMF for 2h. Dead cells were labeled with AnnV/PI and counted 4 h after AMF exposure by confocal microscopy. Results are the mean ± SEM of 4 separate experiments.

## CONCLUSION

In conclusion, unprotected NF@PEG rapidly degraded, released Fe ions and engender ROS upon AMF exposure *in vitro* in an acidic lysosome-like medium, whereas the protected NF@SiO_2_@PEG did not. Both IONP were efficient for MILH, as they both induce similar cell mortality rates (35-40%) with minute amounts of nanoparticles (3 pg γ-Fe_2_O_3_ per cell) and without a perceptible temperature rise during AMF application. Moreover, this study elucidates the origin of iron ions involved in ROS production during MILH by demonstrating that MILH engenders ROS production through Fenton reaction using endogenous Fe^2+^ rather than the Fe^3+^ released from IONPs degradation thanks to local heat at the IONPs surface that is capable to activate and upregulate the Fenton reaction using endogenous Fe^2+^. Moreover, the present work demonstrates that local heat stress generated at the lysosomal level promotes ferroptosis.

## MATERIALS AND METHODS

### Chemical materials

Iron (II) chloride tetrahydrate (FeCl2·4H2O, 98%), iron (III) chloride hexahydrate (FeCl3·6H2O, >97%), iron (III) nitrate nonahydrate (Fe(NO3)3·9H2O, >98%), diethylene Glycol (DEG, 99%), N-methyldiethanolamine (NMDEA, 99%), sodium hydroxide micropellets (NaOH, 98%), nitric acid fuming (HNO3, 69%), hydrochloric acid (HCl, 37%), ethanol (EtOH, anhydrous, ≥99.5%), acetone, ethyl acetate, tetraethyl orthosilicate (TEOS, 99.999%), N-[3-(trimethoxysilyl)propyl]ethylenediamine (EDPS, 97%), ammonium hydroxide solution (30%, ACS reagent), glycerol (BioXtra, ≥99%), dichloromethane (DCM, anhydrous, ≥99.8%), trifluoroacetic acid (TFA, 99% reagent grade), pyridine (anhydrous, 99.8%), N,N⍰.-dicyclohexylcarbodiimide (DCC, ≥99%), petroleum ether (puriss., high boiling, bp 50-70 °C), poly(ethylene glycol) (Mn=2 kDa), sodium borohydride (NaBH4, 99.99%), methanol (MeOH, anhydrous, 99.8%), triethylamine (Et3N, ≥99.5%), thionyl chloride (SOCl2, ≥99%), ethylenediamine (puriss. ≥99.5% (GC)), citric Acid (≥99.5%, ACS reagent), sodium Chloride (≥99.0%, ACS reagent), calcium chloride dehydrate (≥99.0%, ACS reagent), sodium tartrate dehydrate (BioXtra, ≥99%), sodium pyruvate (≥99.0%, ReagentPlus®), sodium lactate (≥99.0%, ReagentPlus®), sodium citrate (≥99.0%), sodium hydrogen phosphate dibasic (99.99%), glycine (≥99.0%, ReagentPlus®), magnesium chloride (anhydrous, ≥98%), sodium sulfate (anhydrous, ≥99.0%) were purchased from Sigma-Aldrich (St Quentin Fallavier, France) and used without further purification. Poly(ethylene glycol) α-ammonium chloride, ω-phosphonic acid (M≈2-3 kDa) was purchased from Specific Polymers, France (https://specificpolymers.com/) and was used without further purification.

### Nanoparticle Synthesis and Functionalization

#### Synthesis of maghemite nanoflowers (NFs)

A mass of 1.082 g (4 mmol) of FeCl_3_·6H_2_O and 0.398 g (2 mmol) of FeCl_2_·4H_2_O is dissolved in 80 g of a liquid mixture of DEG and NMDEA with 1:1 (v/v) ratio (solution A). The resulting solution was then flushed with inert gas (N_2_ or Ar) under stirring for 1 h. In parallel, 0.64 g (16 mmol) of NaOH was dissolved in 40 g solution of polyol 1:1 (v/v) in an ultrasound bath and flushed with an inert gas under stirring for 1h (solution B). Then, solution B was added to solution A and the resulting mixture was flushed with inert gas for 15 min and proceeded to heating with a ramp of 2°C·min^−1^ up to 220°C using an electronically controlled Digi-Mantle heater (Electrothermal™ OMCA0250) under mechanical stirring. After 4h, the resulting black suspension of magnetite Fe_3_O_4_ NPs was removed from the heating mantle to proceed with washing and oxidation. After cooling down to room temperature, the suspension was settled on large ferrite magnets (152×101×25.4 mm^3^, Calamit Magneti™, Milano-Barcelona-Paris) in a beaker for 10 min. After removing all the polyol supernatant by aspiration, a 1:1 mixture of ethyl acetate and ethanol (v/v) was used to wash the solid three times to remove any organic layer covering the nanoparticles originating from polyol decomposition. Next, 8.25 g of ferric nitrate was dissolved in 20 mL of water and boiled before adding to the dispersion of nanoparticles. The resulting suspension was heated to 80°C for 45 min to achieve complete oxidation of the nanoparticles (color shifts from black to brown). The suspension was decanted on magnets to isolate the nanoparticles from the solution. Once aspirated, another 40 mL of a 10 wt% HNO_3_ (2 M) solution was added and the resulting suspension was stirred for 10 min. After magnetic sedimentation, the suspension was washed twice with acetone and twice with diethyl ether. A final aspiration was done and then 20-30 mL of deionized water was added to redisperse the NFs in aqueous medium.

#### Direct grafting of PEG α-ammonium chloride, ω-phosphonic acid onto NFs

For the direct PO-PEG-2k-NH_2_ grafting of NFs, 7 mL of NFs dispersed in water at 16 g□L^−1^ γ-Fe_2_ O_3_ were added in a 20 mL flask. Then, a quantity corresponding to 23.3 PEG macromolecules□nm^−2^ was added to 2 mL of mQ H_2_O and placed in the ultrasonic bath. After 10 min, the PEG solution was added to the NFs under stirring. The pH of the dispersion was set to 2 with HNO_3_ 10 mM. The dispersion is left under agitation overnight. After 12 h, the NPs are washed through tangential ultrafiltration (Merck Millipore, Mw cutoff = 300 kDa) with mQ H_2_O. The washings were stopped once a dilution factor of 10^7^ was reached.

#### Silica coating of NFs

In a 50 mL Erlenmeyer flask containing a volume of 9 mL of an NF dispersion in water (16 g·L^−1^ in γ-Fe_2_ O_3_, developed surface = 10.2 m^2^), a volume of 21 mL of a citric acid solution (14 g·L^−1^) was added. After 5 minutes of stirring, the flocculated nanoparticles were magnetically decanted and washed twice with mQ H_2_O. Finally, the citrate-modified NFs were redispersed in 9 mL of ultrapure water and peptized by adding 10 μL of ammonia (NH_4_ OH at 30% v/v). Then, the aqueous dispersion of citrated NFs (16 g·L^−1^ in γ-Fe_2_ O_3_) was added to a Stöber solution containing 167 mL of ethanol, 44.1 mL of mQ H_2_O and 3.35 mL of NH_4_OH (30% v/v). The stable dispersion was placed in an ultrasonic bath for ~10 min to help break the remaining aggregates. The quantity of TEOS feed was calculated by taking into account the desired SiO_2_ thickness, using the equation below:

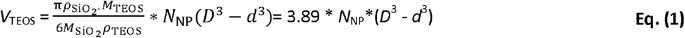

where 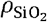 and *ρ* _TEOS_ represent the densities of silica and TEOS (g·cm^−3^) respectively; 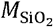 and *M*_TEOS_, the molar masses of silica and TEOS; N_NP_ the number of NPs, D the final diameter, and d the initial diameter of NFs. To generate a 3 nm SiO_2_ layer on N=1.1×10^16^ NFs with a physical diameter of 17.2 nm, a quantity of 319 µL of TEOS was added. The modified NPs were named NF@SiO_2_ at the end of this step.

#### Surface modification of NF@SiO_2_ with an aminosilane coupling agent

NF@SiO_2_ NPs were added to a 50:50 mixture of EtOH/H_2_O so to have a final developed surface of 300 m^2^·L^−1^. Then, an excess of 150 µmol·m^−2^ of EDPS was rapidly injected under vigorous stirring. The NPs quickly flocculate after the EDPS injection. The dispersion was left under stirring overnight. After 12 h, 10 mL of glycerol was added under stirring. The NPs readily disperse upon the addition of glycerol. EtOH and H_2_O were completely removed by heating at 70 °C under vacuum. The colloidal dispersion in glycerol was left at 100 °C under vacuum for 2 h to condense the SiO_2_ coating. The NPs were then transferred through tangential ultrafiltration in MeOH (Merck Millipore, Polyethersulfone PES membrane, Mw cut off = 300 kDa). A few drops of acetic acid were added to the NPs to promote their well-dispersion. Then, washings through tangential ultrafiltration were continued in MeOH until reaching a dilution factor of 10^7^. The NPs were finally stored in anhydrous MeOH and named NF@SiO_2_-NH_2_.

#### Covalent grafting of PEG on NF@SiO_2_-NH_2_

The modification of PEG-2k end chains was inspired by a metal-free Pfitzner-Moffat oxidation step and the protocol was reproduced without further modifications from the work of Adumeau et al. ^37^ Briefly, PEG 2k (20 g, 10 mmol, 1 eq.) was dissolved in 65 mL of anhydrous DMSO. Traces of water were removed by evaporating 20 mL of DMSO at 80°C under vacuum. After cooling to room temperature, 50 mL of anhydrous DCM, followed by pyridine (2.01 mL, 25 mmol, 2 eq.) and TFA (960 µL, 12.5 mmol, 1 eq.) were then added under stirring at 4°C for 30 min. Then, DCC (15.5 g, 75 mmol, 6 eq.) was added. The mixture was kept for 20 h at 4°C under stirring in the dark. The white precipitate corresponding to dicyclohexylurea formed as a by-product of DDC, was separated by filtration and washed with 30 mL of cold DCM. The remaining solution was then added to a cold mixture of petroleum ether and propan-2-ol to induce the precipitation of PEG-2k-aldehyde. After filtration, the PEG-2k-al was solubilized in 50 mL of DCM and then filtered to remove traces of dicyclohexylurea. The remaining solution was then added to 500 mL of diethyl ether, placed at 4°C to precipitate the PEG-2k-al, and then filtered under vacuum. The washing process was repeated 3 times. The final precipitate was dried under vacuum and stored at −20°C. The PEG-2k-al (41 µmol, 81 mg) was then solubilized in anhydrous MeOH and kept under stirring for 20 min. The NPs dispersion in MeOH (5.6 m^2^ of NF@SiO_2_-NH_2_) was added dropwise to the PEG solution while stirring vigorously. After 5 min, 22.4 µmol of trimethylamine (Et_3_N) was directly injected and the dispersion was left stirring for 10 min. Then, half of the MeOH was eliminated under vacuum. The final dispersion, containing the NPs and PEG in 5.86 mL of anhydrous MeOH, was then heated at 70°C under reflux overnight. The next morning, NaBH_4_ (1 mmol) was added and left under stirring for 10 min before introducing 1 mL of HCl 1M to neutralize the dispersion. The PEG excess was removed through tangential ultrafiltration in MeOH (Merck Millipore, PES membrane, Mw cut off = 300 kDa). In a third step, the terminations of PEGylated NPs were aminated. To do so, in a flask, a volume of 1.42 mL of DMF is added to 8.2 mL of PEGylated NF@SiO_2_ (4.6 g.L^−1^ in γ-Fe_2_ O_3_) dispersed in MeOH, placed in a 5 mL round bottom flask. Then, MeOH was eliminated by heating the dispersion at 70 °C under vacuum. Water traces were also removed by increasing the heat to 90 °C under vacuum for 5 min. The dispersion was then cooled down in an ice bath. A volume of 20 μL of SOCl_2_ (0.27 mmol) and 11 μL (0.14 mmol) of pyridine was added to the dispersion under vigorous stirring. After 5 minutes, the dispersion is placed at 70 °C for 2 h. After the reaction, the NPs in DMF were mixed in a 500 mL beaker with 300 mL of MeOH. The dispersion was then transferred in the ultrafiltration cell (Merck Millipore, PES membrane, Mw cut off = 300 kDa) to pursue with two washings to remove excess SOCl_2_. The NPs were finally dispersed in MeOH and added to 1.4 mL of anhydrous DMF. The MeOH was eliminated under vacuum at 70 °C. Under magnetic stirring, a volume of 18 μL of ethylenediamine (0.27 mmol) was added to the dispersion of NPs and placed at 70 °C overnight. After the reaction, the dispersion was mixed in 300 mL MeOH and transferred into the tangential ultrafiltration cell (Merck Millipore, PES membrane, Mw cut off = 300 kDa) to proceed with washings in EtOH (twice) and then H_2_O. The NPs were finally dispersed in H_2_O and named NF@SiO_2_-PEG.

#### Conjugation of NF@PEG and NF@SiO_2_@PEG with fluorophore and gastrin

The NF@PEG and NF@SiO_2_@PEG nanoparticles were then decorated with ~20 molecules of the fluorescent label NHSDY647-PEG1 (Dyomics GmbH, Jena, Germany) and ~100 molecules of a synthetic replicate of gastrin (Cys-Lys-Ser-Ser-Glu-Ala-Tyr-Gly-Trp-Nle-Asp-Phe-NH2, Covalab), and termed respectively as NF@PEG@gastrin and NF@SiO_2_@PEG@Gastrin. An aliquot of nanoparticles was sonicated for 5 min on melting ice (Bioblock Scientific 88154). Firstly, the linkage of the fluorophore NHS-DY647-P1 or NHS-DY549-P1 (20 molecules per IONP) was performed in phosphate buffer pH 8.3, at room temperature for 1 h. Then, 3-Maleimidopropionic acid N-hydroxysuccinimide ester (BMPS, 1n NH2) in solution in DMF was added and incubated at room temperature for 1h. After 5 washes with 10 volumes of H_2_O, gastrin (~100 peptides per nanoparticles) in solution in 30% DMF, 70% H_2_O was added in phosphate buffer pH 7.2 and let to react for 2h. Finally, free maleimide functions were saturated by addition of an excess of cysteine (10n NH2) in phosphate buffer pH7.2. After 8 washes with 10 volumes of H_2_O, NF@PEG@gastrin and NF@SiO_2_@PEG@Gastrin were stored at 4°C.

### Physico-chemical characterization of nanoparticles

Transmission Electron Microscopy (TEM) samples were prepared by depositing a drop (10 µL) of nanoparticles suspension on carbon coated copper grids (Lacey/thin double carbon film Cu-300LD, 300 mesh, Pacific Grid Tech, San Francisco, CA). The excess of the droplet was aspired with a paper filter to leave a thin liquid film on the TEM grid. TEM images were obtained using a Jeol JEM-1400+ instrument operated at 120 kV, and digital micrographs were obtained with a Smart Orius 1000 Gatan camera. NP size distribution and diameter were obtained by manually measuring 300 NPs of each batch with ImageJ software (https://imagej.nih.gov/ij/). Size-histograms were fitted to a normal distribution law via Origin software.

Dynamic light scattering (DLS) operated in backscattering mode i.e. at 165° angle (Vasco™ Flex, Cordouan Technologies™, Pessac, France) was used to calculate the hydrodynamic intensity-average size and polydispersity index (PDI) as determined with the 2nd order cumulant analysis. In practice, five runs of 40 s duration were acquired, the Z-average diameter (Zave) and PDI were averaged, and a standard deviation was calculated. Dispersions of NF@PEG@gastrin and NF@SiO_2_@PEG@Gastrin at 0.1 mg/ml were prepared in phosphate buffer pH 7.4 before measurements. The suspension was on ice for 10 min. After temperature equilibrium to 20°C, particle hydrodynamic size was measured.

The ζ potential measurement was recorded on WALLIS™ analyzer based on Laser Doppler Electrophoresis (LDE) measurement technology (Cordouan Technologies™, Pessac, France). Before the measurement, the NPs were dispersed in a 1 mM HEPES buffer (pH=7.4) and placed in the dip cell composed of amorphous carbon electrodes. The measurement was repeated 5 times for each batch.

The Thermo-gravimetric analysis (TGA) experiments were conducted on a TGA-Q50 from TA instruments. For this, 5-10 mg of nanoparticles were first dried at 90 °C under vacuum for 12 h. Then, the obtained powder was placed in a platinum crucible and heated from room temperature to 600 °C with a rate of 10 °C·min-1 under N2 flux.

Magnetization experiments under DC field were conducted on a Vibrating Sample Magnetometer (VSM) Microsense EZ-7. The magnetization was recorded under a range of applied magnetic field intensities H from 0 to 4000 kA·m^−1^ with regularly spaced data points. The magnetization curves were fitted with Langevin law convolved with a Log-normal distribution of diameters:

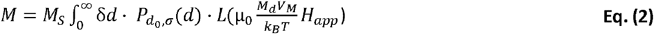

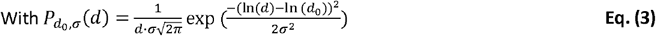

where *µ*_0_ is the vacuum magnetic permeability, *H*_*app*_ the intensity of the applied field, *M*_*d*_ the magnetic domain magnetization, 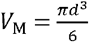 the magnetic core volume for a spherical particle of diameter *d*, T being the absolute temperature and *k*_*B*_ the Boltzmann constant. Also, 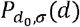, corresponds to the nanoparticle size distribution fitted by log-normal law where *d*_*0*_ is the median value of the distribution and *σ* the standard width of the log values. The integral on diameters *d* is calculated numerically for a finite increment *δd* = 0.1 nm and only *d*_*0*_ and *σ* as adjustable parameters (the domain magnetization being simply *M*_*d*_ = *M*_*s*_ · *ρ*, where *M*_*S*_ is the saturation magnetization per mass of iron oxide and is *ρ* = 4800 kg·*m*^−3^ the mass density of the solid (taken as the tabulated value for maghemite). From the Langevin fits on the experimental data of *M(H)* and the specific relations of the log-normal law, we obtain the mean number-averaged magnetic size of the NPs:

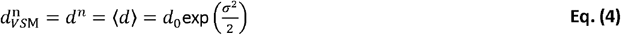

with its standard deviation:

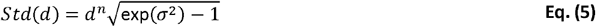

The size distribution obtained by the Langevin fit can also be weighted by the volume fraction of the NPs expressed *Phi*(*d*) to obtain the volume or weight-averaged magnetic size:

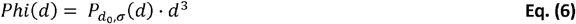

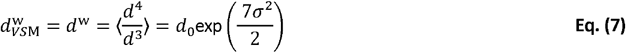

Dynamic hysteresis loops were measured by AC magnetometry with a pick-up coil technology ^38^ using the AC Hyster™ setup commercialized by NanoTech Solutions company (Ntsol, Madrid, Spain). An aliquot of 40 µL suspension with a concentration of 3 g·L^−1^ (containing a dry γ-Fe_2_ O_3_ mass of ≈ 1.2 × 10^−4^ g) was put at the bottom of a 3 mm diameter 4 inches length NMR tube (VWR, France). Then the magnetization cycles *M(H*) were measured thrice (waiting for 45 s between each measurement for the sample to cool down) at a series of magnetic field amplitudes (*H*_*max*_) ranging from 4 to 24 kA⋅m^−1^ and at a frequency (*f*) varied in this series: 146, 217, 280 and 344 kHz. The three measured cycles were averaged and normalized by the exact weight of iron oxide known from iron titration to compute the mass magnetization in A·m^2^·kg^−1^. To cope with unavoidable drifts of the pick-up coil calibration, each curve *M(H)* was adjusted by a normalization factor so that the mass magnetization measured under an AMF of amplitude *H*_*max*_ =24 kA⋅m^−1^ was identical to the value measured by *VS*M magnetometry under a DC magnetic field of same intensity *H*_DC_ =24 kA⋅m^−1^.

For specific absorption rate measurements, a 300 μL solution of NF@PEG and NF@SiO_2_@PEG at 7 mg γ-Fe_2_O_3_/mL was used. The samples were placed in a magnetic inductor (Fives Celes, Lautenback, France) generating a 93 kHz alternating magnetic field with an amplitude of 0 to 38 kA⋅m^−1^. The temperature increases was measured during 100 s. A blank sample containing water was used in parallel in order to remove the contribution of eddy current to the temperature increase. Temperature measurements were performed using a thermal probe (Reflex, Neoptix, Canada). The specific absorption rate was calculated using standard formula and taking into account the specific heat of water.

### Analysis of *in vitro* NF degradation

A volume of 1.3 mL of NPs dispersion (concentration varied from 0.5, 3, and 7 g·L^−1^ γ-Fe_2_ O_3_) in ALF was added in a 1.5 mL conical Eppendorf® tube. The cap of the tube was pierced on its center to immerse the optical fiber temperature probe (OpSens) in the sample. The temperature probe was placed in the center of the Eppendorf® tube and taped on top of its cap to keep it in place throughout the experiment. The tube was then positioned under the coil of the DM3 (nB nanoScale Biomagnetics, Zaragoza, Spain). An identical tube was placed under the other coil of the device to follow the dissolution kinetics by withdrawing 200 μL every 20 min for 2 h. The temperature of the sample was first regulated at 37°C thanks to the circulation of hot water in rubber pipes. The samples were also covered with cloth to limit heat dissipation in the environment. The magnetic field was turned on once the temperature of the sample is stable at 37.0 ± 1.5 °C and left on for 2h. To check the reproducibility of the AMF set-up and the used conditions, the experiment was repeated 3 times for each type of NPs for concentrations of 0.5 and 3 g·L^−1^ γ-Fe_2_O_3_ and twice for 7 g·L^−1^ γ-Fe_2_O_3_.

Two sets of experiments in water bath, acting as the “control” of the degradation assays under AMF were conducted: 1/ Each set of NPs (NF@PEG and NF@SiO_2_@PEG) dispersed in ALF with final concentrations of 0.5, 3 and 7 g·L^−1^ γ-Fe O were incubated for 2h at 37 °C. Then, the NPs were centrifuged at high speed (60×10^3^ g, 25 min, 10°C). The supernatant was collected and its absorbance was recorded at 350 nm to check the degradation of the NPs; 2/ Each set of NPs (NF@PEG and NF@SiO_2_@PEG) was incubated in a water bath with a temperature set at the value reached during AMF through magnetic hyperthermia, corresponding to 45 and 65°C for each type of NPs at a concentration of 0.5 and 3 g·L^−1^ γ-Fe_2_ O_3_ respectively.

### *In vitro* ROS production

16 μg of NF@PEG@Gastrin or NF@SiO_2_@PEG@Gastrin were incubated into 84 μL of 0.2 M acetate buffer pH 4.5, composition) and exposed or not to alternating magnetic field (AMF: 21 kA.m^−1^, 275 kHz) delivered by a magnetic inductor for 2 h. Then 1 μL of of 3,3⍰,5,5⍰-Tetramethylbenzidine (TMB) (20 mg/mL in DMSO) and 5 μL of H2O2 (10 mM) were added. The UV− vis absorbance spectra of oxidized TMB was recorded via a microplate reader (SpectroStar BMG) for 45 minutes.

### Cell Culture

Pancreatic cancer cell line MiaPaca2 was permanently transfected with the human CCK2 receptor (CCK2R) using the cDNA encoding CCK2R subcloned in pcDNA3 vector (BD Biosciences Clontech) and Lipofectamine 2000 (Invitrogen), selected with 600 µg/ml geneticin (Sigma-Aldrich) and termed MiaPaca2-CCK2. MiaPaca2 and MiaPaca2-CCK2 were cultured in complete DMEM medium (DMEM GlutaMAX™, Thermofisher Scientific) containing 10% FBS, 100 IU/ml penicillin/streptomycin (Penicillin-Streptomycin P4333; Sigma-Aldrich) in a humidified atmosphere at 95% air and 5% CO2 at 37°C. MiaPaca2-CCK2 cells were grown with 600 µg/ml geneticin (Sigma-Aldrich).

### Analysis of MNP cytotoxicity

10^4^ cells/well of MiaPaca2-CCK2 was seeded in a 96-well plate, grown overnight and treated with increased concentrations of NF@PEG@Gastrin or NF@SiO_2_@PEG@Gastrin in complete DMEM medium for 24, 48 or 72h. Ten microliters of MTT (3-(4,5-dimethylthiazolyl-2)-2,5-diphenyltetrazolium bromide) at 5 mg/mL were added and incubated for 2 h at 37 °C. After cell culture medium removal, 100 µL of dimethylsulfoxid were added and incubated for 1 h at 37 °C, and the absorbance values were measured at 570 nm. The cells maintained in the incubation medium without MNPs served as negative controls.

### Hemolysis Assay

Heparinized blood was obtained from male CD-1 mice and male Sprague-Dawley rats by cardiac puncture under anesthesia, and from male human healthy volunteers (with no major medical history and absence of medications). Red blood cells were incubated with increasing concentrations of NF@PEG@Gastrin or NF@SiO_2_@PEG@Gastrin, 1 mg/ml Triton X-100 (positive control) or PBS (negative control) for 4h at 37C. The samples were centrifuged at 1,000 g for 10 min. Optical density of the supernatant was measured at 540 nm with a Tecan Safire 2 plate reader. All measurements were performed in triplicates. The % is calculated relative to the positive control (Triton X-100) inducing 100% hemolysis. Results are expressed as mean ± sem.

### Analysis of MNP uptake by Flow Cytometry

10^5^ MiaPaca2 or MiaPaca2-CCK2 cells were seeded onto 24-well plates and grown overnight. For the kinetic, cells were incubated with NF@PEG@Gastrin or NF@SiO_2_@PEG@Gastrin ([γ-Fe_2_O_3_] = 16 µg/mL) for 2 to 72h at 37°C in DMEM medium. For the specificity, cells were incubated with increased concentrations of NF@PEG@Gastrin or NF@SiO_2_@PEG@Gastrin ([γ-Fe_2_O_3_] = 2 to 64 µg/mL) for 72h. Then, cells are washed 3 times with phosphate-buffered saline (PBS) containing 0.5% BSA (Bovine serum albumin), collected with 300 µL of trypsin and centrifuge 10 min at 1500 rpm. The cells were resuspended in 300 µL of PBS, 2 mM EDTA. Cell-associated fluorescence was determined using a BD FACSCalibur™ flow cytometer.

### Quantification of MNP uptake by relaxometry

~10^6^ MiaPaca2-CCK2 cells were seeded onto 35 mm dishes in complete DMEM medium. Cells were incubated with NF@PEG@Gastrin or NF@SiO_2_@PEG@Gastrin ([γ-Fe_2_O_3_] = 16 µg/mL) in complete DMEM medium for 72 h, at 37°C in a 5% CO2 atmosphere, washed twice with ice-cold PBS, and centrifuge 10 min at 1500 rpm.

### MNP localization by confocal microscopy

50 × 10^3^ cells/compartment were plated onto 4-compartment Cellview culture dishes (Greiner Bio-One). After overnight growth, cells were incubated with NF@PEG@Gastrin or NF@SiO_2_@PEG@Gastrin ([γ-Fe_2_O_3_] = 16 µg/mL) in complete DMEM medium for 72 h. For lysosome staining, cells were incubated for 15 minutes in the presence of 10 nM LysoTracker Green DND-26 (excitation: 488 nm, Life technologies). MNP (excitation: 633 nm) and Lysotraker co-localization was analyzed using a LSM780 confocal microscope (Zeiss).

### MNP localization by transmission electron microscopy

10^5^ MiaPaca2-CCK2 cells were grown in 6-dishes plates (Thermo Scientific™) and incubated with NF@PEG@Gastrin or NF@SiO_2_@PEG@Gastrin ([γ-Fe_2_O_3_] = 16 µg/mL) for 72 h. After removing medium, cells were fixed with 4% glutaraldehyde in Sorensen buffer for 4 h at 4°C. Fixed cells were washed with cold Sorensen buffer for 12 h and post-fixed in 1% osmium tetra oxide (osmium 2%, saccharose 0.25 M, Sorensen 0.05 M) for 1 h at 20°C, followed by washings with distilled water and uranyl acetate 2% for 12 h at 4°C. The cells were dehydrated by sequential washings of 10 min in 30, 50, 70 and 95% ethanol followed by 3 washings of 15 min in absolute ethanol. Cells were then embedded in EMBed 812 resin for 12 h. Resin polymerization was obtained at 60°C for 48 h. Ultrathin sections of 70 nm were prepared, stained with uranyl acetate and lead citrate and examined with a transmission electron microscope (Hitachi HU12A, Japan) operating at 75 kV.

### Local heat quantification using molecular thermometers

25 × 10^3^ cells/cm^2^ MiaPaca2-CCK2 cells were seeded onto 35-mm dishes. After overnight growth, cells were incubated with NF@PEG@Gastrin or NF@SiO_2_@PEG@Gastrin ([γ-Fe_2_O_3_] = 16 µg/mL) in complete DMEM medium for 72 h, washed and exposed to AMF using a miniaturized home-made electromagnet placed in a CELLView dish, ^23,39^. This permits to follow in real-time the evolution of the fluorescence of the MNPs under the application of AMF and thus to have an access to the local temperature. Before and during AMF application, fluorescence intensities of DY549 (excitation: 540 nm) was measured from confocal microscopy images LSM780 confocal microscope, Zeiss) using ImageJ software. 20–30 cells/experiment were analyzed from 4 independent experiments.

### Cell Exposure to alternating magnetic field

Cells were seeded 24 h before the experiments onto four-compartment Cellview dishes (Greiner Bio-One), grown overnight and incubated with NF@PEG@Gastrin or NF@SiO_2_@PEG@Gastrin for 72h at 37 °C in complete DMEM medium to allow MNP internalization and accumulation in lysosomes. The incubation medium was removed and cells were rinsed with complete medium. Finally, cells were exposed or not to alternating magnetic field (AMF: 21 kA.m^−1^, 275 kHz) delivered by a magnetic inductor for 2 h. The temperature of the incubation medium was strictly maintained at 37°C and controlled using a thermal probe (Reflex, Neoptix, Quebec City, QC, Canada) placed in the incubation medium. At the end of the experiments, cells were placed in a humidified atmosphere at 5% CO2 at 37°C for further analysis.

### Analysis of cell Death

25 × 10^3^ cells/compartment of MiaPaca2-CCK2 cells were seeded onto four-compartment Cellview dishes, incubated with NF@PEG@Gastrin or NF@SiO_2_@PEG@Gastrin in complete DMEM medium for 72h, washed and exposed to AMF (21 kA.m^−1^, 275 kHz) as previously described. 4 h after AMF exposure, dead cells were labeled with FITC-annexin V (AnnV) and/or propidium iodide (PI) (Cell Meter Annexin V apoptosis assay kit, AAT Bioquest, Sunnyvale, CA, USA) in accordance with the manufacturer’s instructions. The counting of labeled cells was carried out through the analysis of confocal microscopy images (LSM510, Zeiss) representing populations of 2000–3000 cells/experiment using Image J software.

### Analysis of cell proliferation

10^4^ cells/compartment of MiaPaca2-CCK2 cells were seeded onto 4-compartment Cellview dishes (Greiner Bio-One), grown overnight and incubated with NF@PEG@Gastrin or NF@SiO_2_@PEG@Gastrin ([γ-Fe_2_O_3_] = 16 µg/mL) for 72h. Then, the cells were washed, incubated in complete DMEM medium and exposed to AMF (21 kA.m^−1^, 275 kHz) for 2h, 3 times every 48 h. The effects of magnetic field treatments were investigated on cell proliferation by counting the cell number by a cell counter (Beckman cell counter z2) once per day for 6 days.

### Analysis of ROS production

25 × 10^3^ cells/compartment of MiaPaca2-CCK2 cells were seeded onto 4-compartment Cellview dishes (Greiner Bio-One), grown overnight, incubated with NF@PEG@Gastrin or NF@SiO_2_@PEG@Gastrin ([γ-Fe_2_O_3_] = 16 µg/mL) for 72 h in complete DMEM medium, washed, then incubated in complete DMEM medium. For experiment with Ferrisatin-II, cells were incubated NF@PEG@Gastrin or NF@SiO_2_@PEG@Gastrin ([γ-Fe_2_O_3_] = 16 µg/mL) for 72 h in complete DMEM medium, washed, then incubated in SVF-free DMEM medium in the presence or absence of 50 µM of Ferristatin-II (Sigma-Aldrich). Cells were exposed to AMF (21 kA.m^−1^, 275 kHz) for 2 h and incubated with CellROX Green reagent (Molecular probes, Excitation wavelegnth: 488nm) immediately after AMF exposure, in incubation medium according to manufacturer’s instructions. The quantification of ROS production was performed by analyzing the intensity of CellROX Green reagent labeling of confocal microscopy images (LSM780 confocal microscope, Zeiss) using Image J software. 2,000-3,000 cells/experiments were analyzed in at least 4 independent experiments.

### Analysis of lysosomal membrane permeabilization

25 × 10^3^ cells/compartment of MiaPaca2-CCK2 cells were seeded onto 4-compartment Cellview dishes (Greiner Bio-One), grown overnight, incubated with NF@PEG@Gastrin or NF@SiO_2_@PEG@Gastrin ([Fe_2_O_3_] = 16 µg/mL) for 72 h in complete DMEM medium, washed, then incubated in complete DMEM medium. Cells were exposed to AMF (21 kA.m^−1^, 275 kHz) for 2 h, incubated with medium containing 10 nM LysoTraker Green DND-26 (Invitrogen, excitation wavelength: 488 nm) for 15 minutes, rinsed with incubation medium. Lysosome integrity was determined by analyzing Lysotraker fluorescence intensity by confocal microscopy images (LSM780 confocal microscope, Zeiss).

### Fe(II) detection

To detect intracellular Fe(II), FerroOrange (Dojindo, Japan) was used based on the manufacturer’ s protocol. Brie fly, the harvested cells were washed with SVF-free DMEM medium and then incubated with 1 µM FerroOrange working solution at 37 ° C for 30 min. Fluorescence images were captured by a LSM780 confocal microscope (Zeiss, excitation: 543 nm). Analysis of Fe(II) was performed by measuring the intensity of fluorescence per cell.

### Statistical Analysis

Results are expressed as the mean ± SEM of at least three independent experiments. The statistical analysis was performed using ANOVAc tests, as mentioned in each figure legend. Differences were considered significant when p < 0.05.

## ACKNOWLEDGMENTS

The authors thank the Genotoul TRI platforms: Cellular Imaging and Flow Cytometry platforms of I2MC/INSERM, and CMEAB (centre de microscopie electronique appliquée à la biologie) for the excellent technical support. We thank Coralie Genevois for technical assistance for AMF experiments at Vivoptic TBMCore platform, UAR CNRS 3427, INSERM US 005, Univ. Bordeaux, France. Vivoptic is a France Life Imaging (FLI) labelled platform. J.JD. was supported by grants from Ligue Nationale Contre le Cancer. This work was funded by the canceropole Grand Sud-Ouest.

---

**Figure S1.**
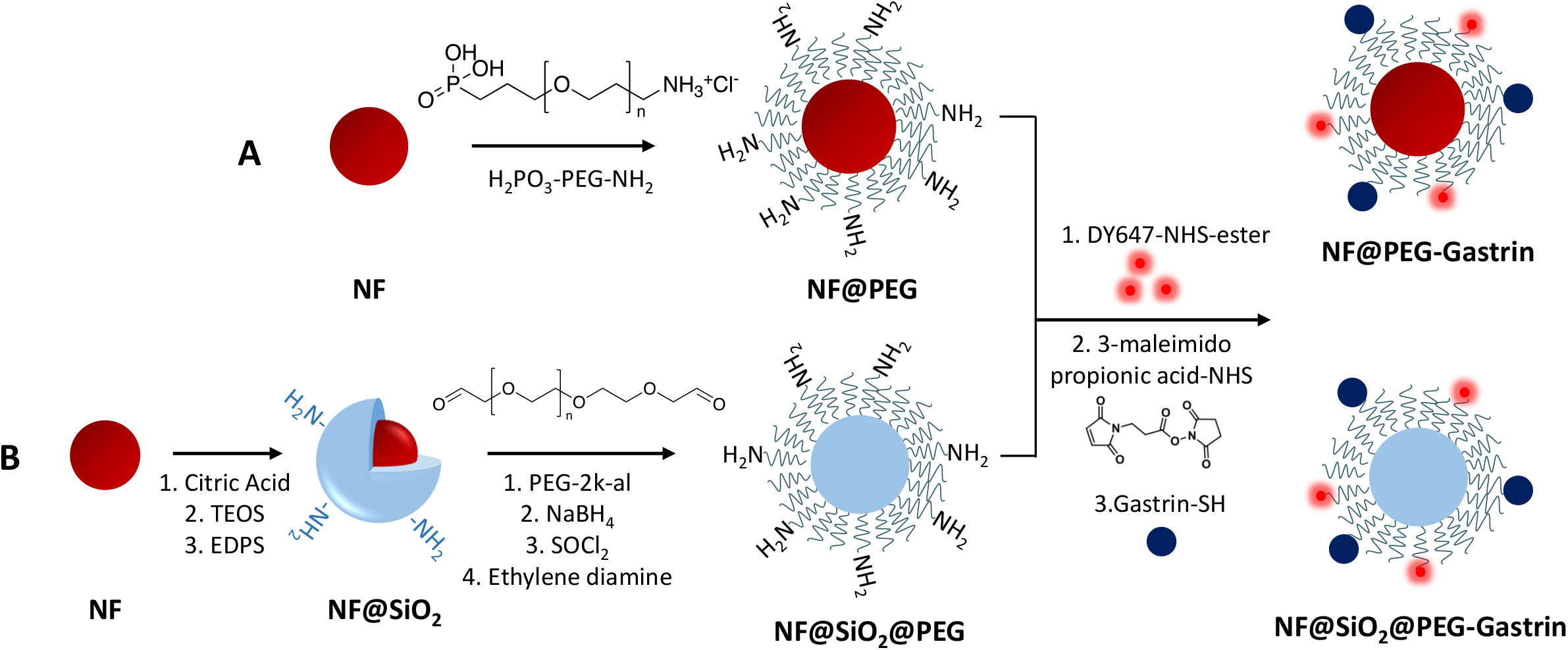

**Figure S2.**
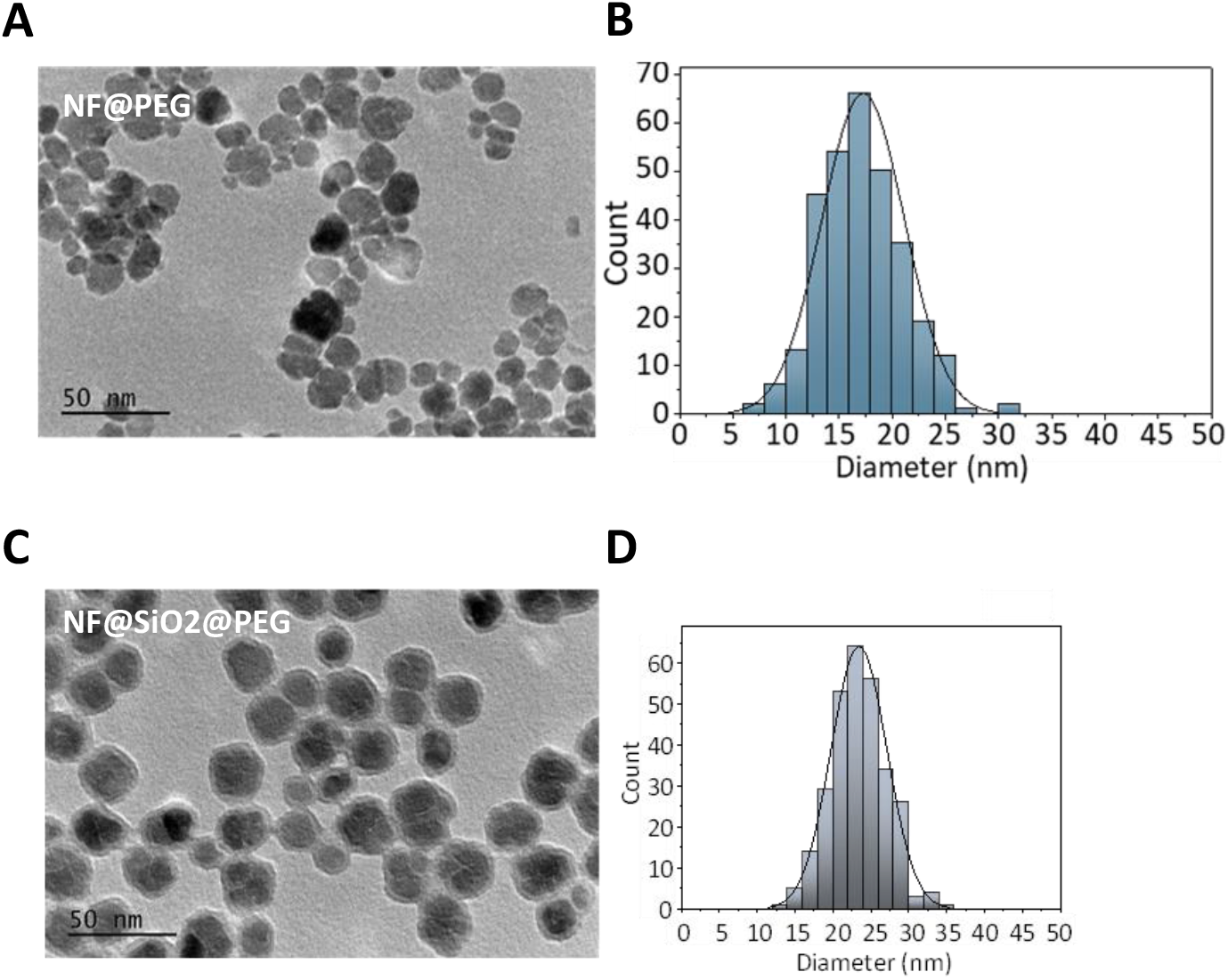

**Figure S3.**
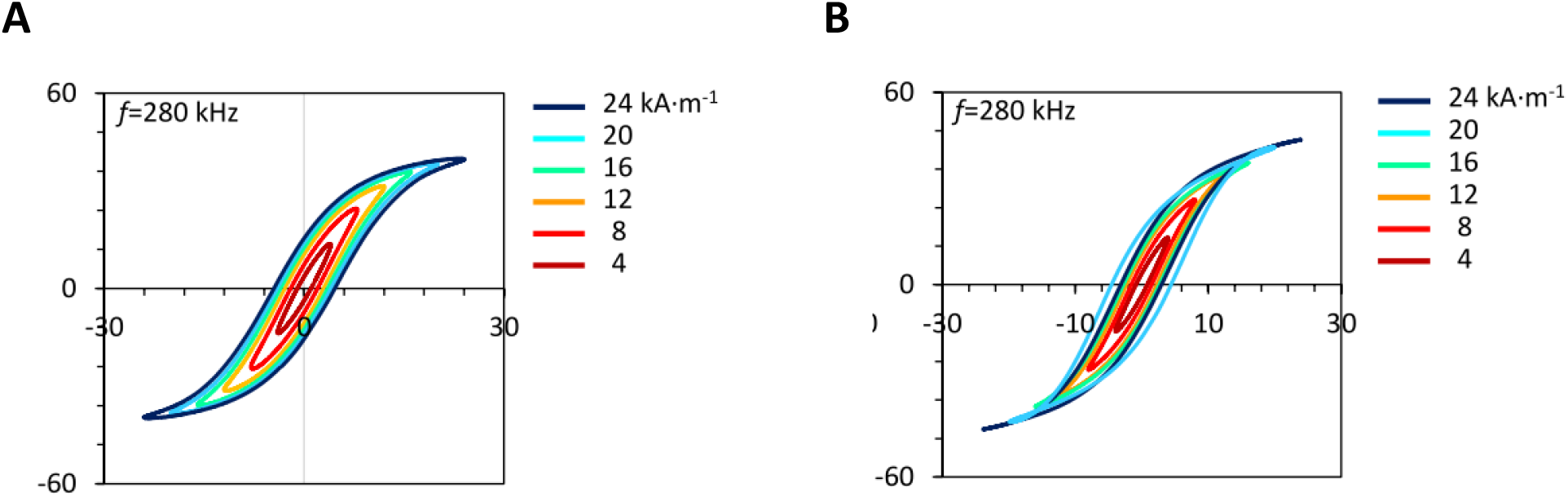

**Figure S4.**
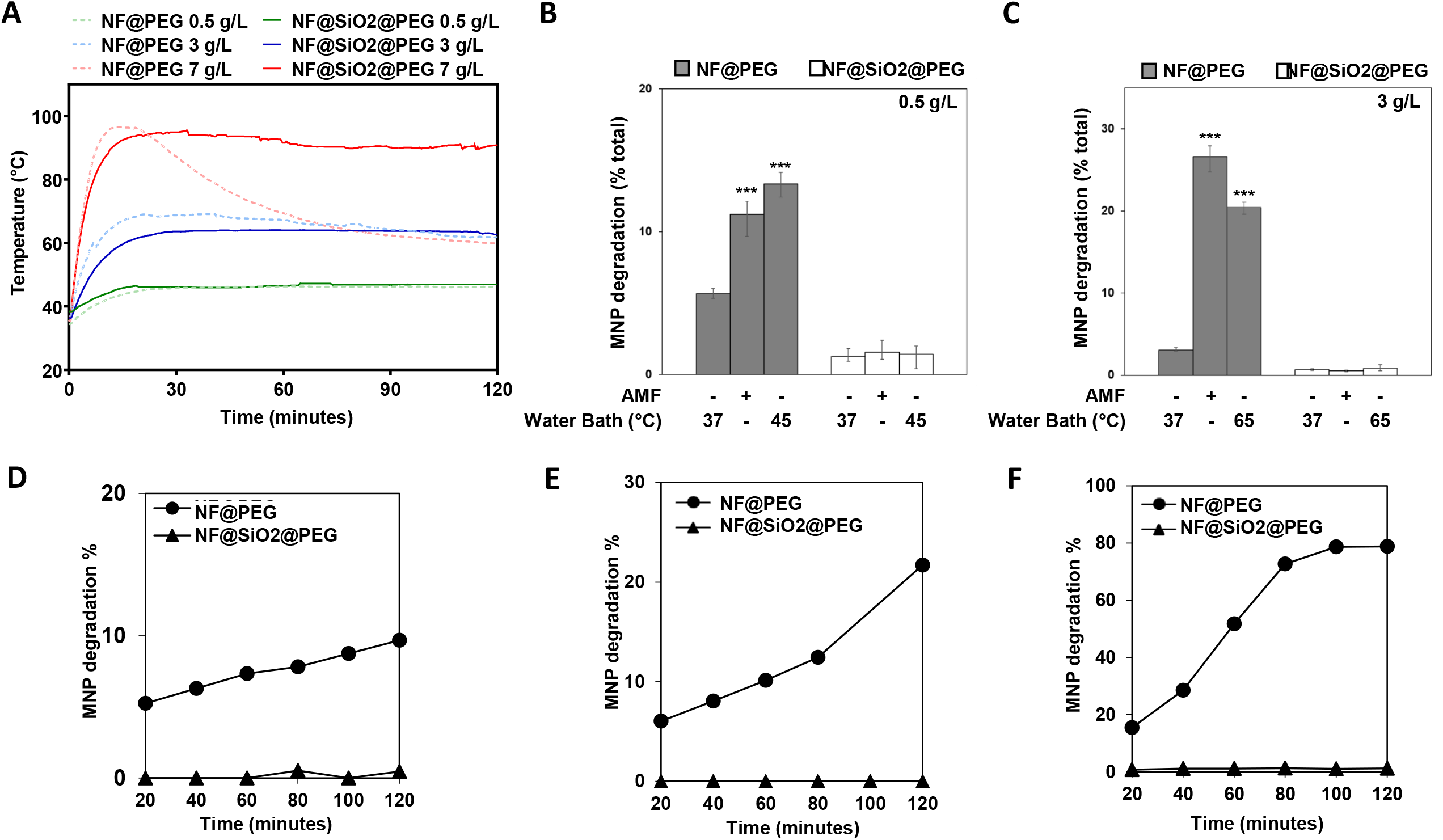

**Figure S5.**
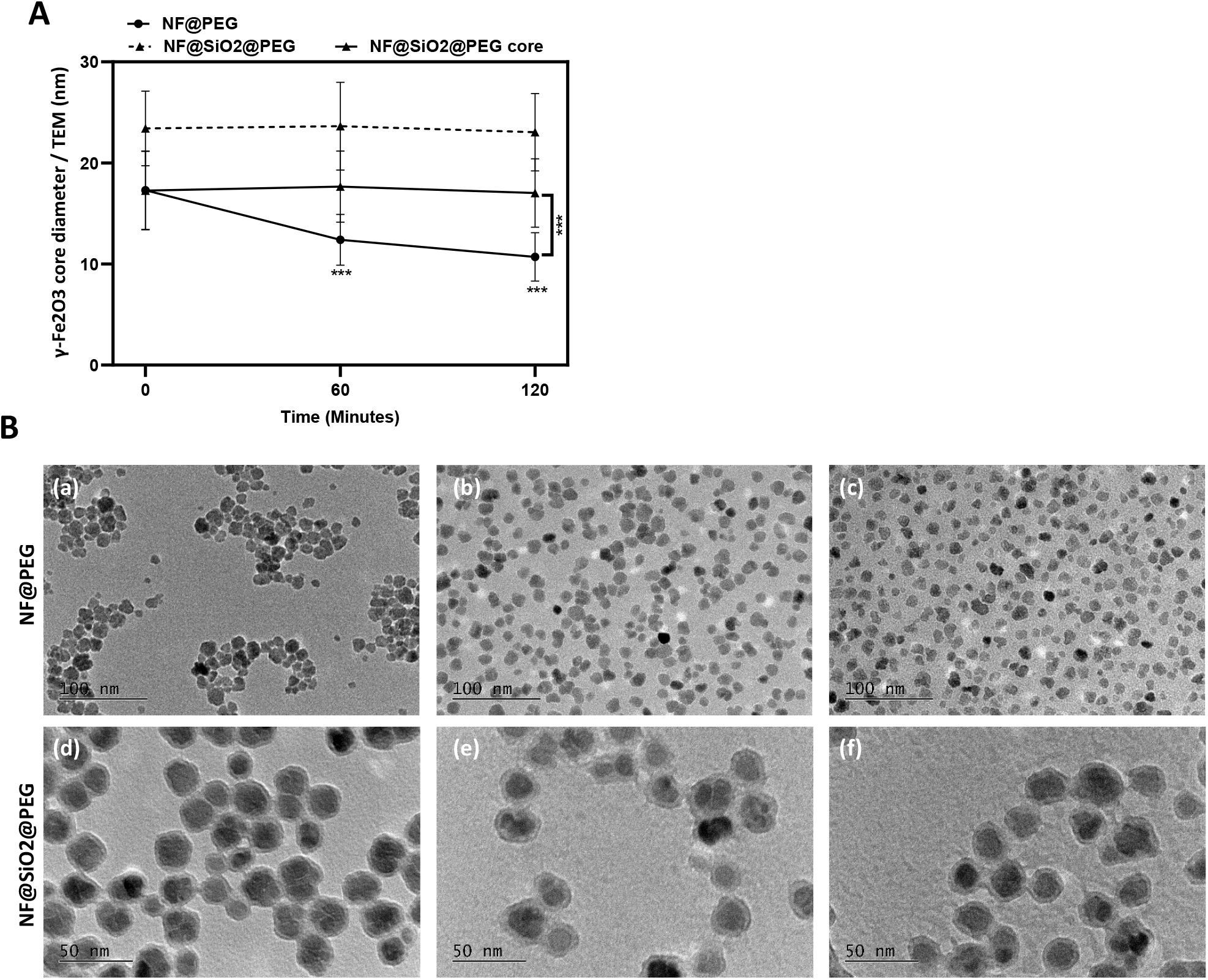

**Figure S6.**
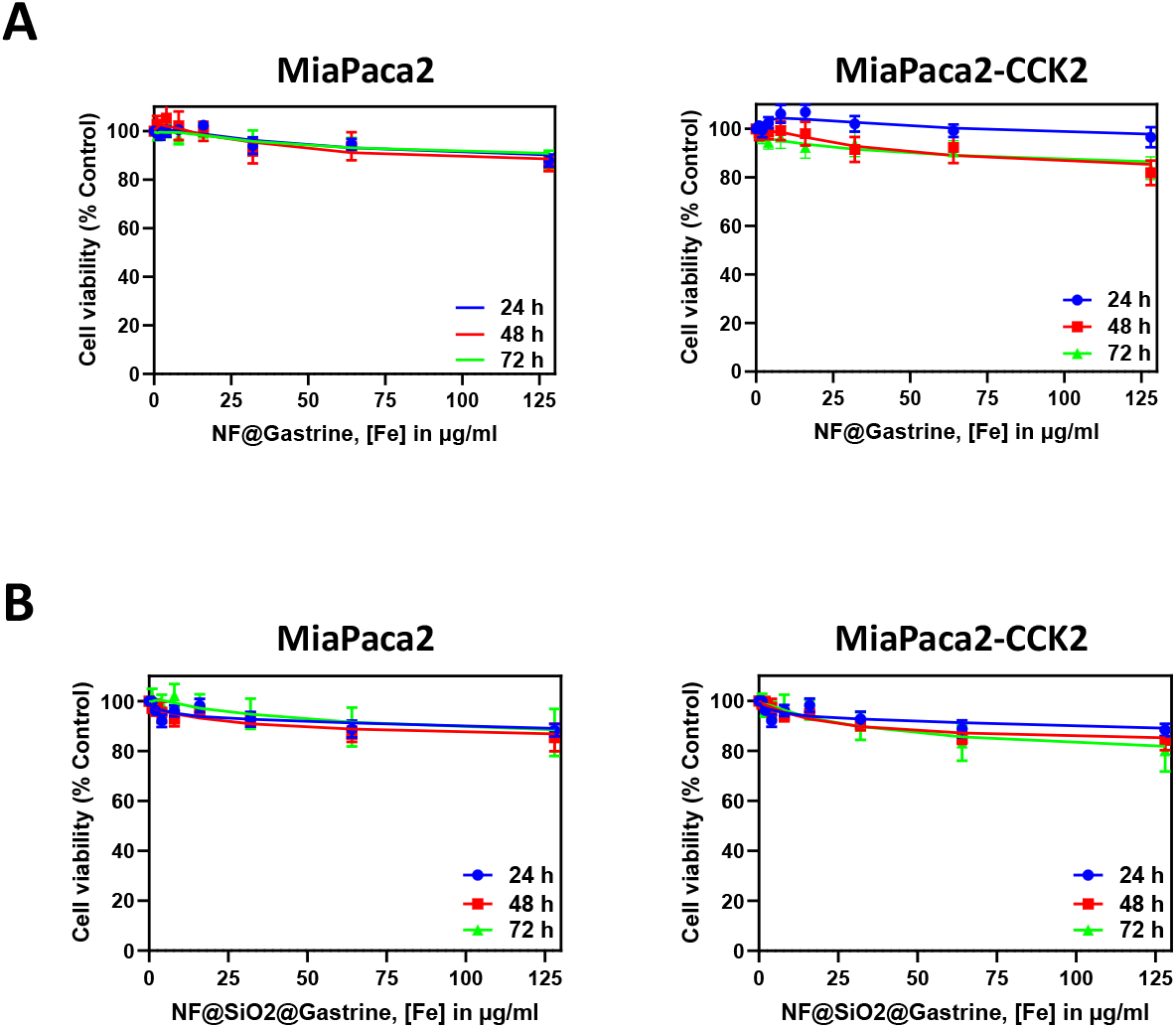

**Figure S7.**
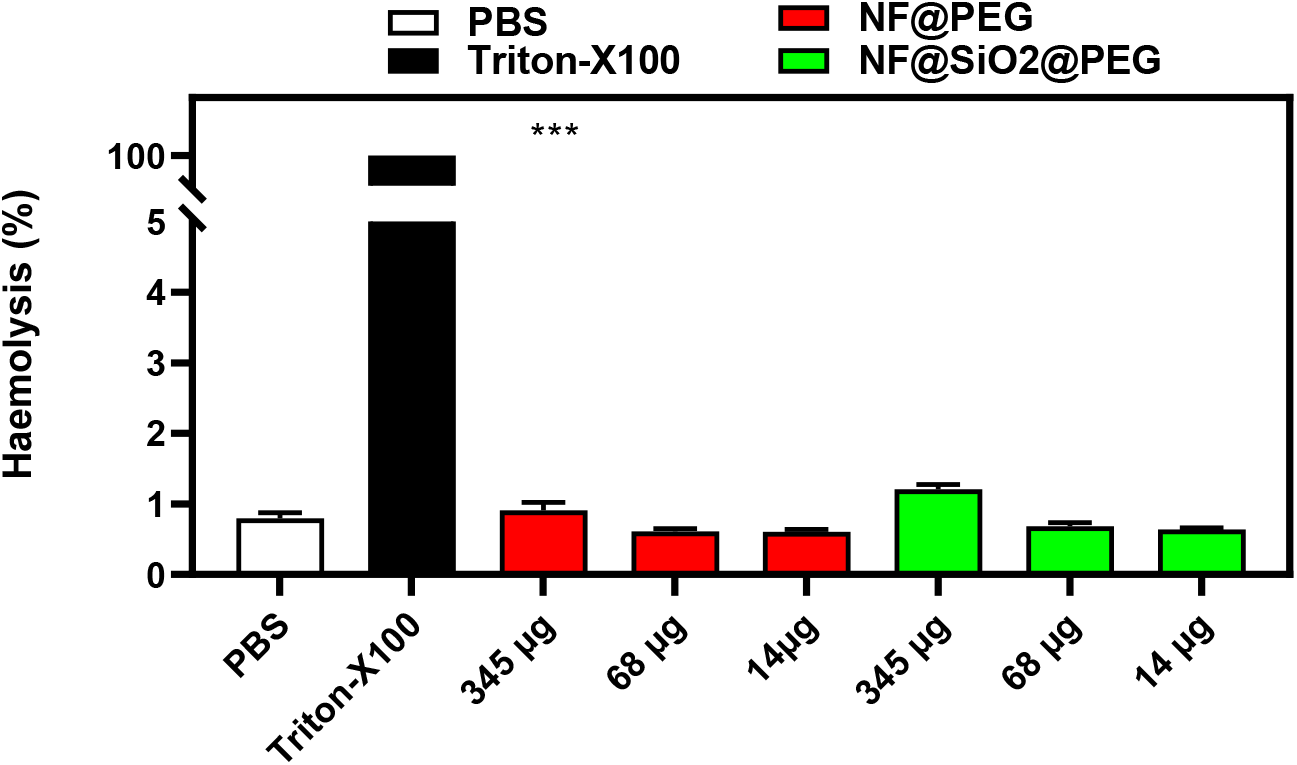

**Figure S8.**
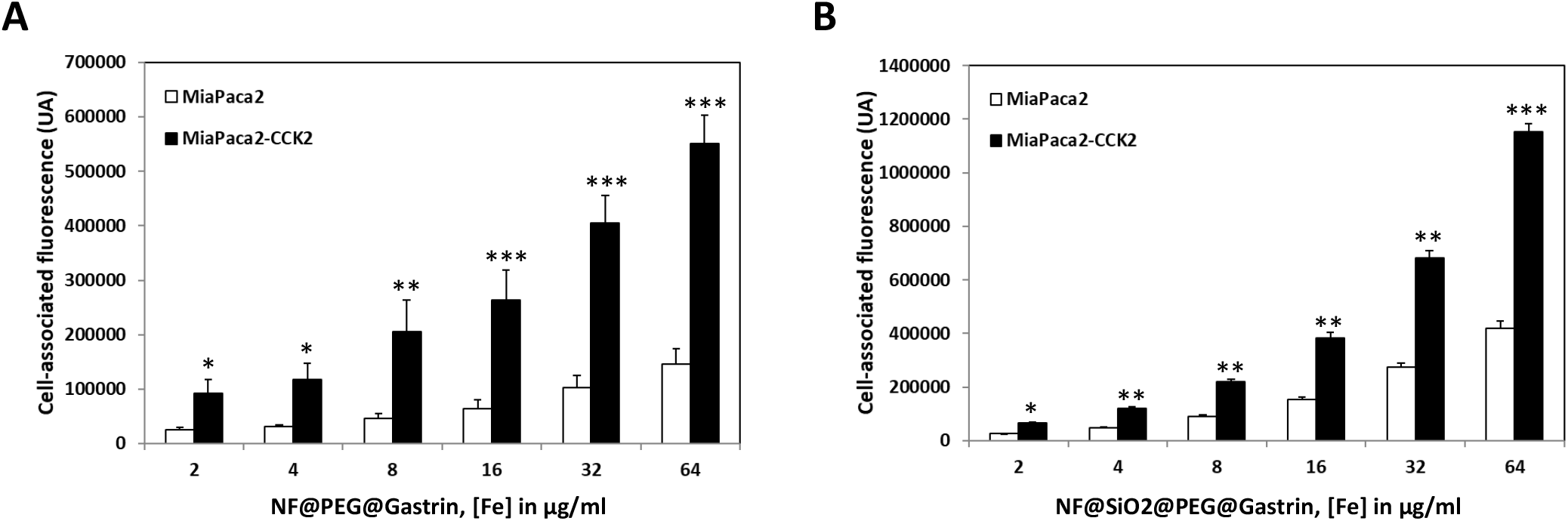

**Figure S9.**
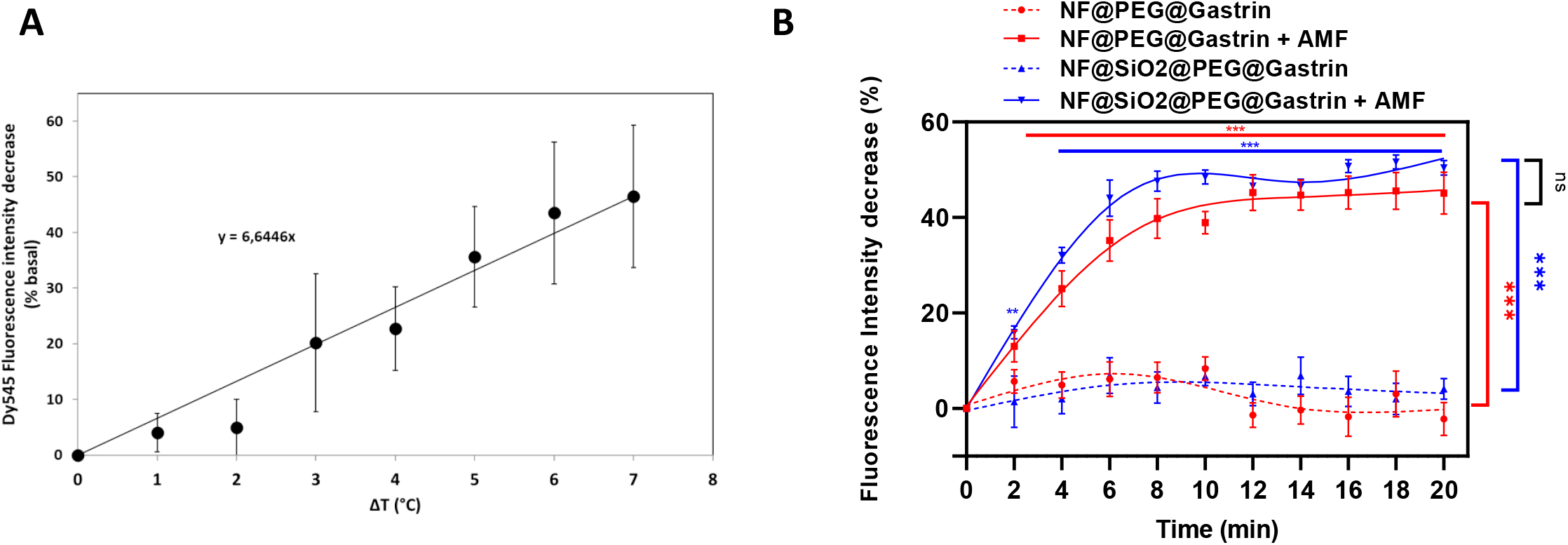

**Figure S10.**
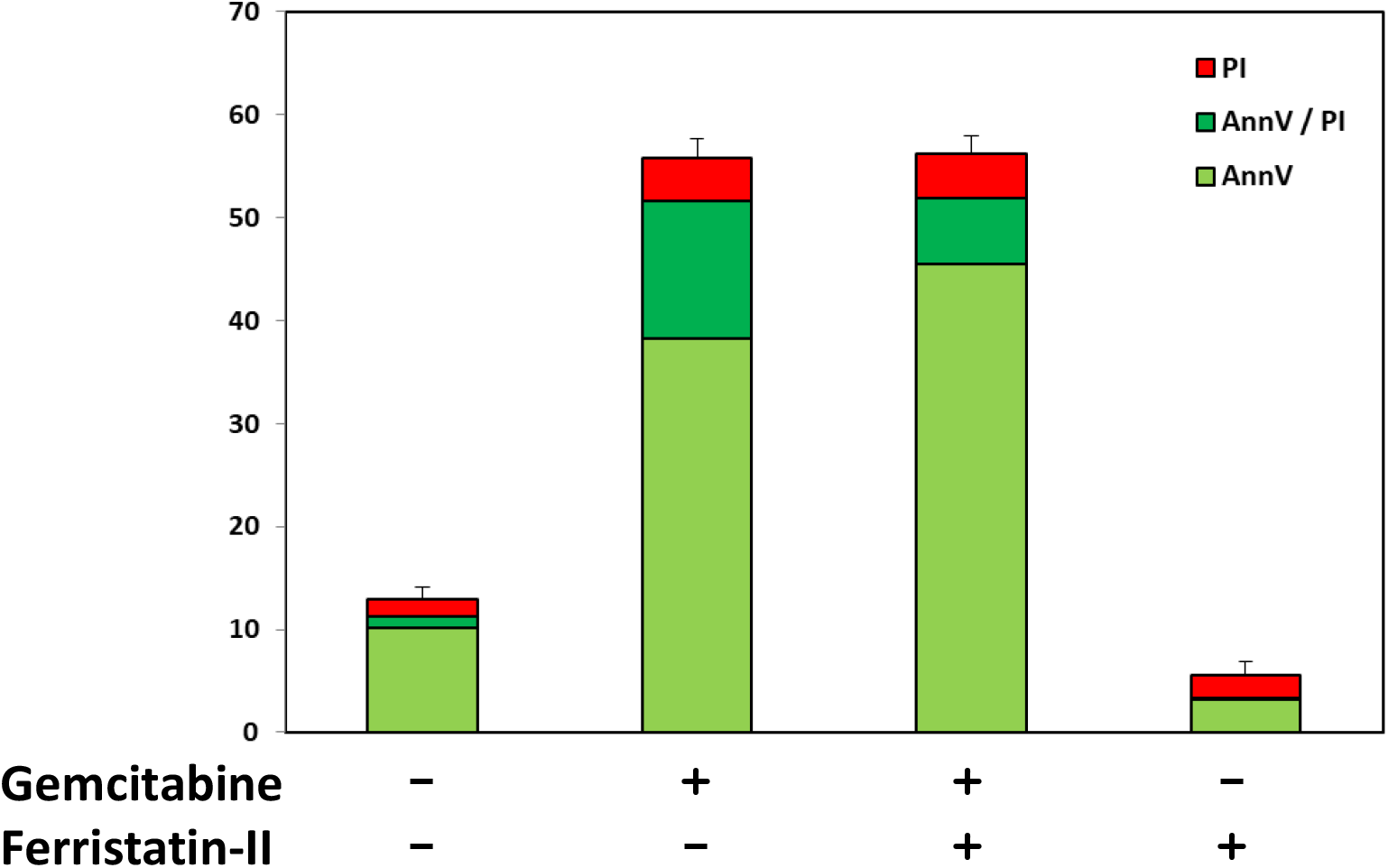

