## supplementary figures for "Origin of Fe ions in ROS production induced in magnetic hyperthermia anti-cancer nanotherapy: release from iron oxide nanoparticles or not?"

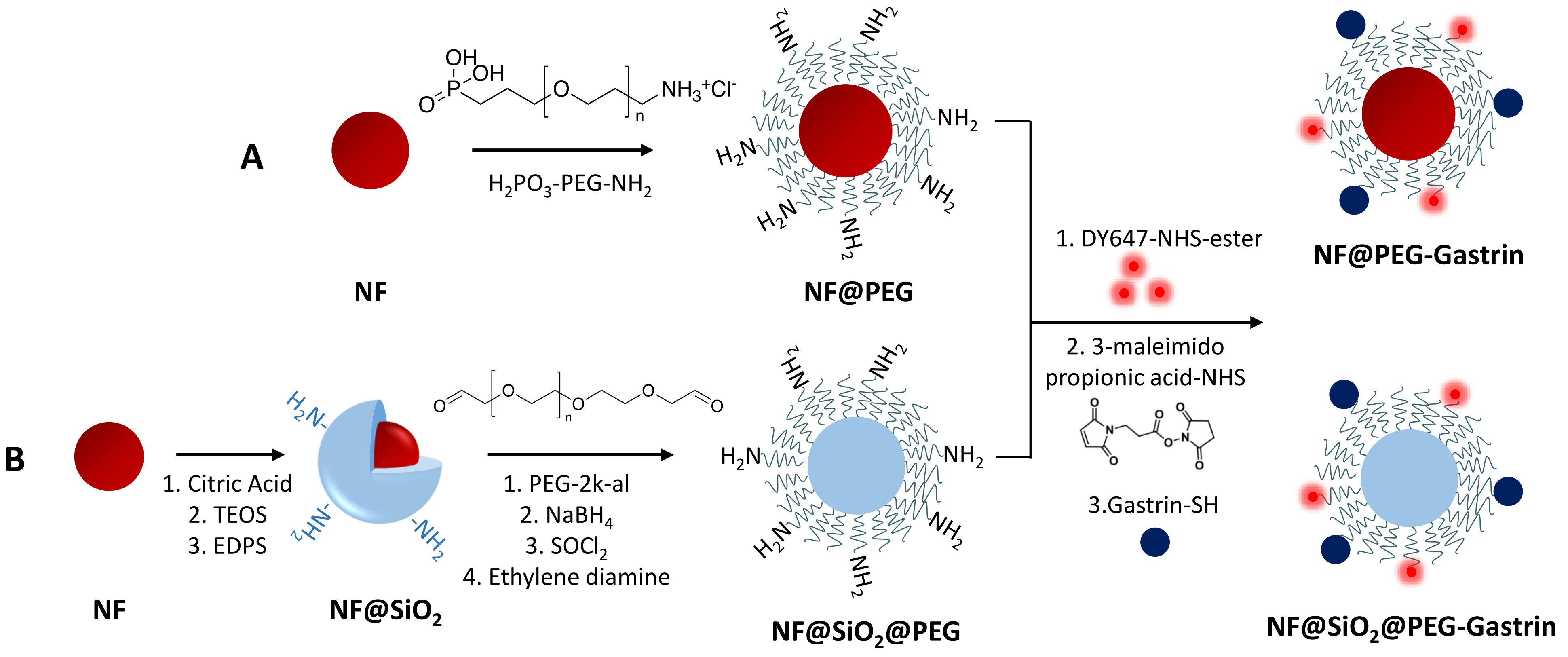
A
B
H2PO3-PEG-NH2
NF
NF@PEG
1. Citric Acid
2. TEOS
3. EDPS
NF@SiO2
1. PEG-2k-al
2. NaBH4
3. SOCl2
4. Ethylene diamine
NF@SiO2@PEG
1. DY647-NHS-ester
2. 3-maleimido propionic acid-NHS
3.Gastrin-SH
NF@PEG-Gastrin
NF@SiO2@PEG-Gastrin

**Figure S1**

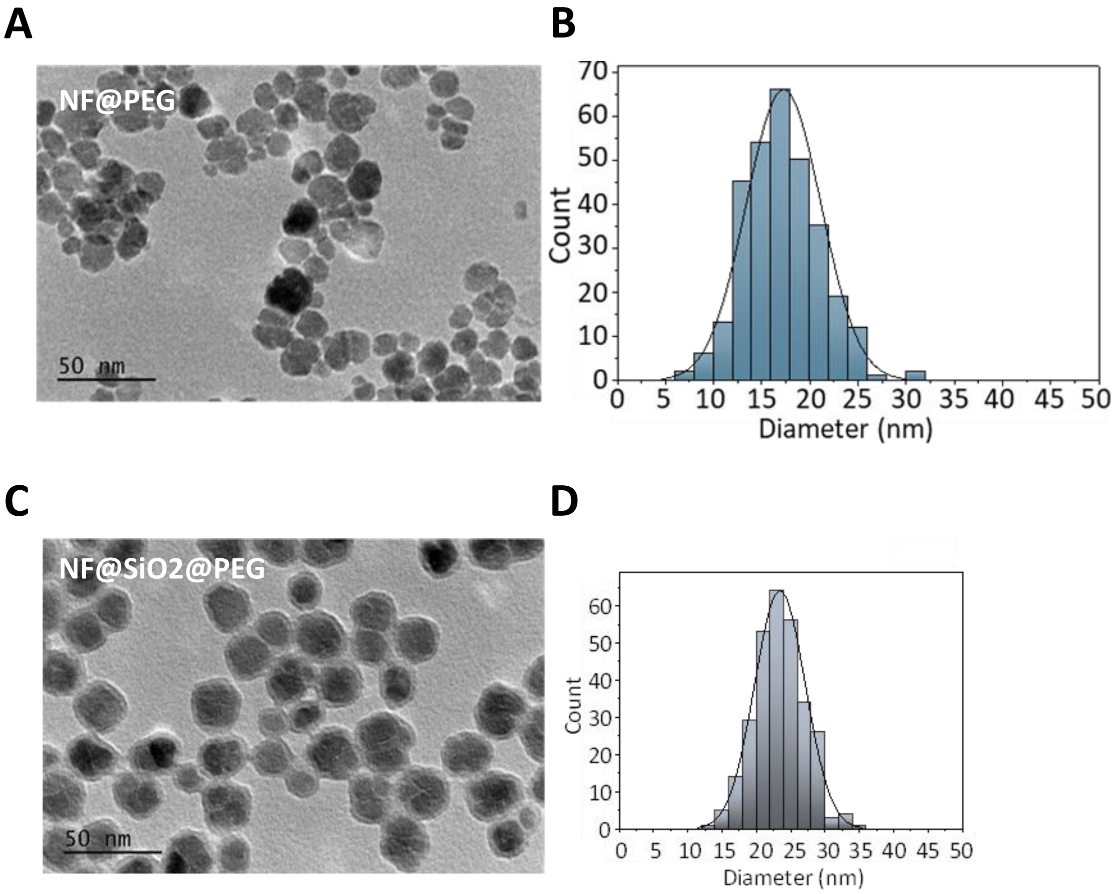
A
B
C
D
NF@PEG
50 nm
Count
Diameter (nm)
0
10
20
30
40
50
60
70
5
15
25
35
45
NF@SiO2@PEG
50 nm
Count
Diameter (nm)

**Figure S2**

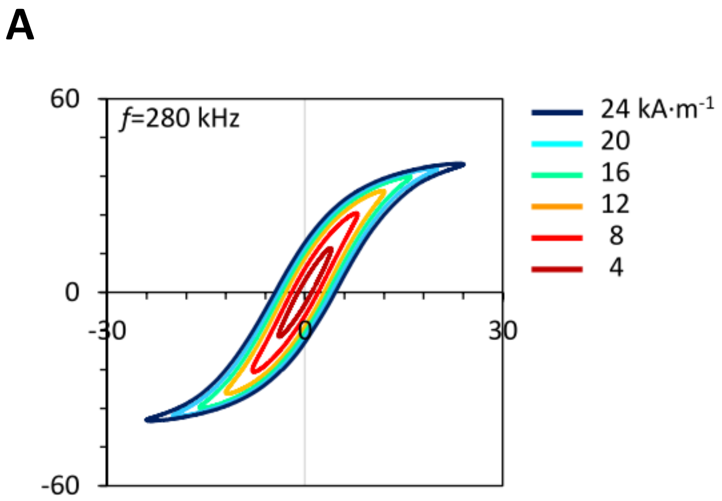
A
60
f=280 kHz
0
-30
0
30
-60
24 kA·m-1
20
16
12
8
4

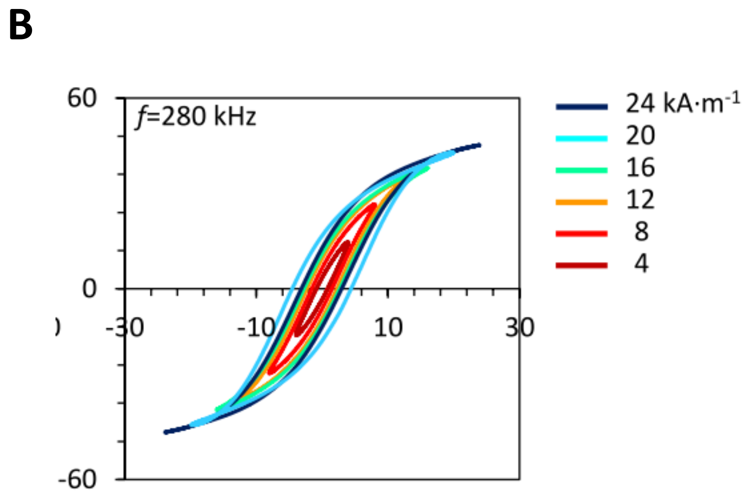
B
60
f=280 kHz
0
-30
-10
10
30
-60
24 kA·m-1
20
16
12
8
4

**Figure S3**

| | NF@PEG | NF@PEG@Gastrin | NF@SiO2@PEG | NF@SiO2@PEG@Gastrin |
|---|---|---|---|---|
| **$D_{TEM}$ (nm)** | 17.3 ± 2.6 | | 23.4 ± 3.7<br>(SiO2 shell 3.1 ± 0.2) | |
| **$D_H$ (nm)** | 50.3 ± 4.0 | 75.8 ± 5.1 | 64.5 ± 6.9 | 79,3 ± 7.3 |
| **PDI** | 0.25 ± 0.15 | 0.15 ± 0.01 | 0.17 ± 0.02 | 0.15 ± 0.03 |
| **ζ (mV)** | +14.0 ± 2.0 | -21,6 ± 1.1 | +22.6 ± 5.7 | -1.0 ± 0.5 |
| **RF/D (PEG regimen)** | 2.4 (brush) | | 2.7 (brush) | |
| **Ms ($kA.m^{-1}$)** | 356 | | 340 | |
| **Dn (nm)** | 9.1 ± 0,8 | | 10.2 ± 0,9 | |
| **Dw (nm)** | 11.2 ± 0,7 | | 11.9 ± 1,1 | |

**Table 1**

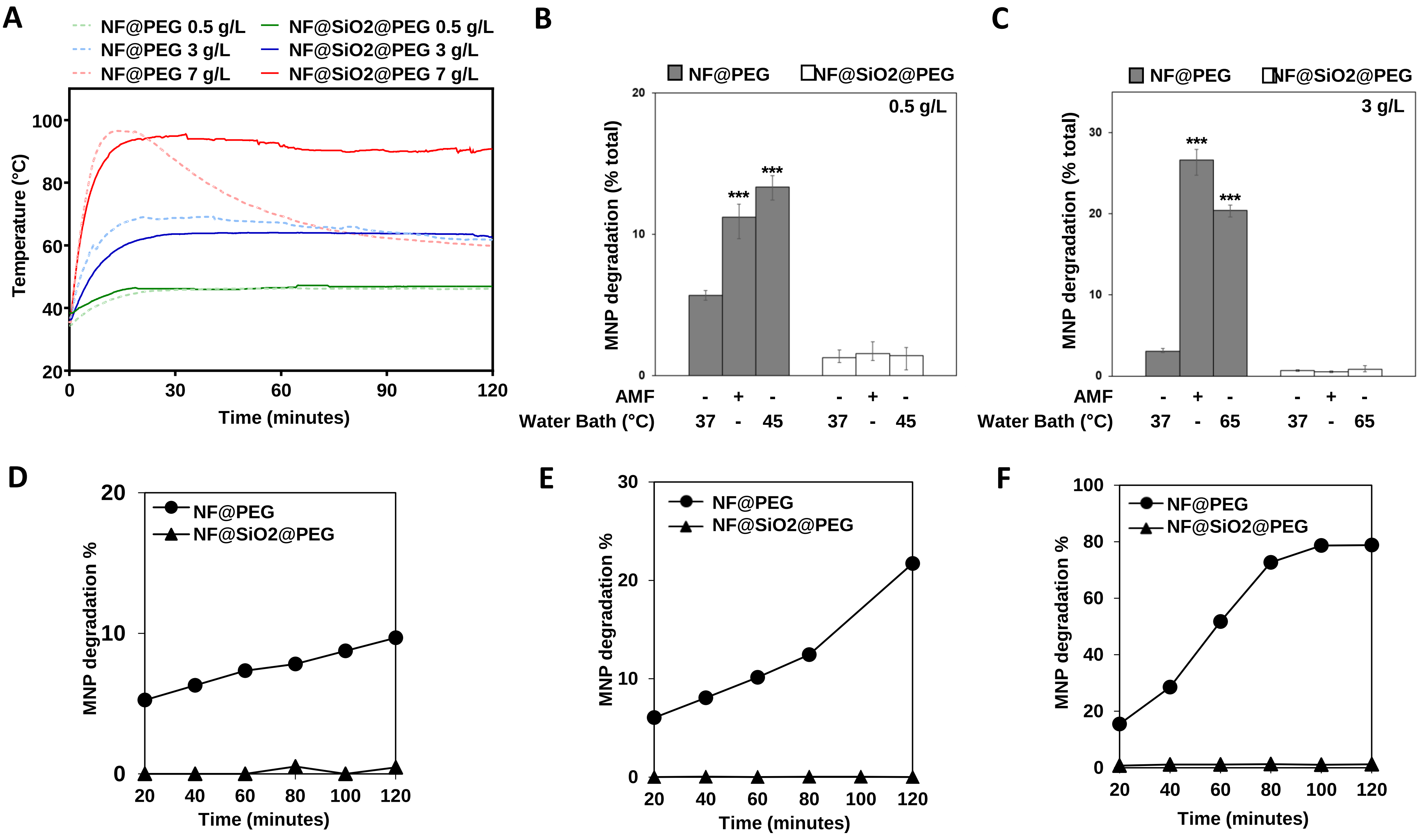

A
NF@PEG 0.5 g/L
NF@PEG 3 g/L
NF@PEG 7 g/L
NF@SiO2@PEG 0.5 g/L
NF@SiO2@PEG 3 g/L
NF@SiO2@PEG 7 g/L
Temperature (°C)
Time (minutes)
B
NF@PEG
NF@SiO2@PEG
0.5 g/L
MNP degradation (% total)
AMF
Water Bath (°C)
C
3 g/L
MNP dergradation (% total)
D
E
F
MNP degradation %


Figure S4

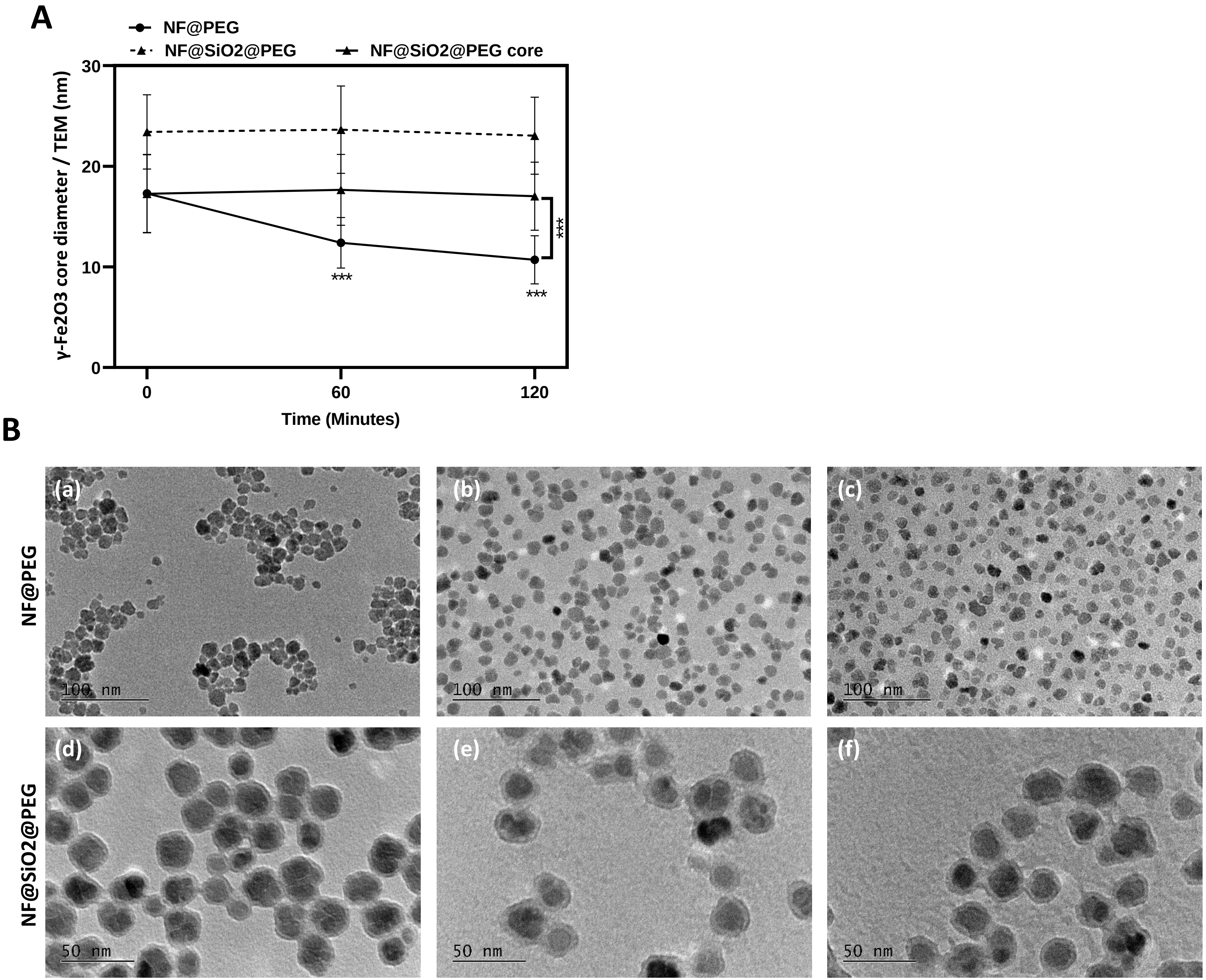
A
NF@PEG
NF@SiO2@PEG
NF@SiO2@PEG core
γ-Fe2O3 core diameter / TEM (nm)
30
20
10
0
***
***
***
0
60
120
Time (Minutes)
B
NF@PEG
NF@SiO2@PEG
(a)
(b)
(c)
(d)
(e)
(f)
100 nm
100 nm
100 nm
50 nm
50 nm
50 nm

Figure S5

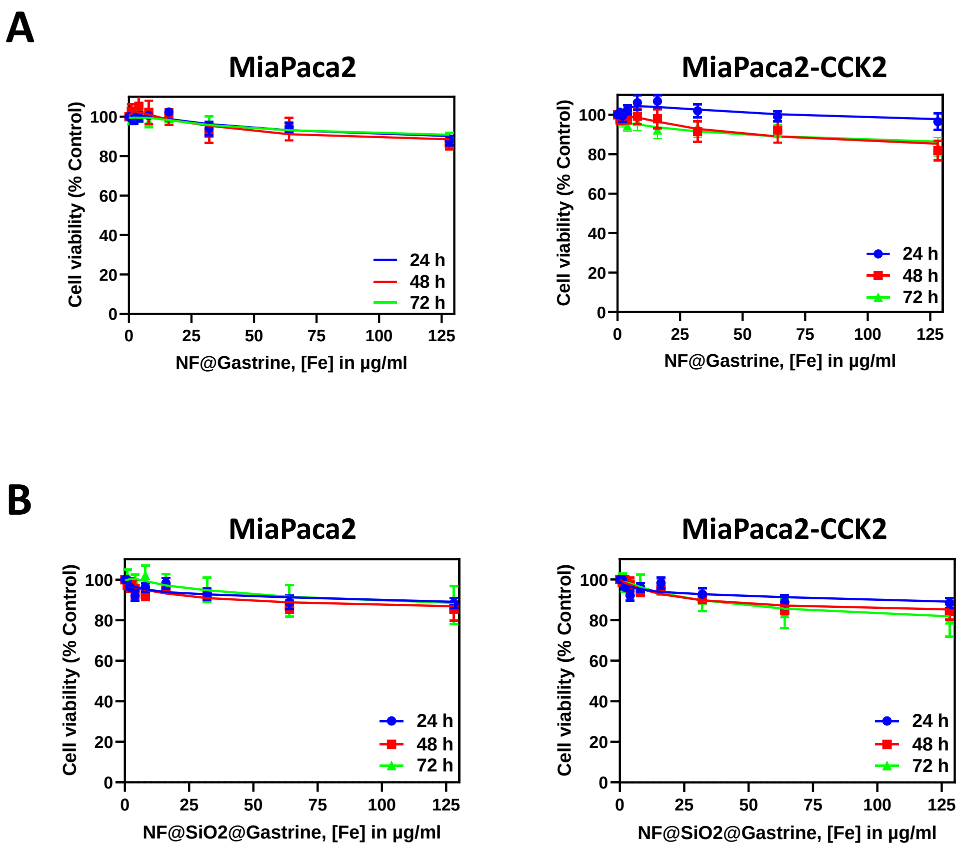
A
MiaPaca2
Cell viability (% Control)
NF@Gastrine, [Fe] in µg/ml
24 h
48 h
72 h
MiaPaca2-CCK2
Cell viability (% Control)
NF@Gastrine, [Fe] in µg/ml
24 h
48 h
72 h
B
MiaPaca2
Cell viability (% Control)
NF@SiO2@Gastrine, [Fe] in µg/ml
24 h
48 h
72 h
MiaPaca2-CCK2
Cell viability (% Control)
NF@SiO2@Gastrine, [Fe] in µg/ml
24 h
48 h
72 h

**Figure S6**

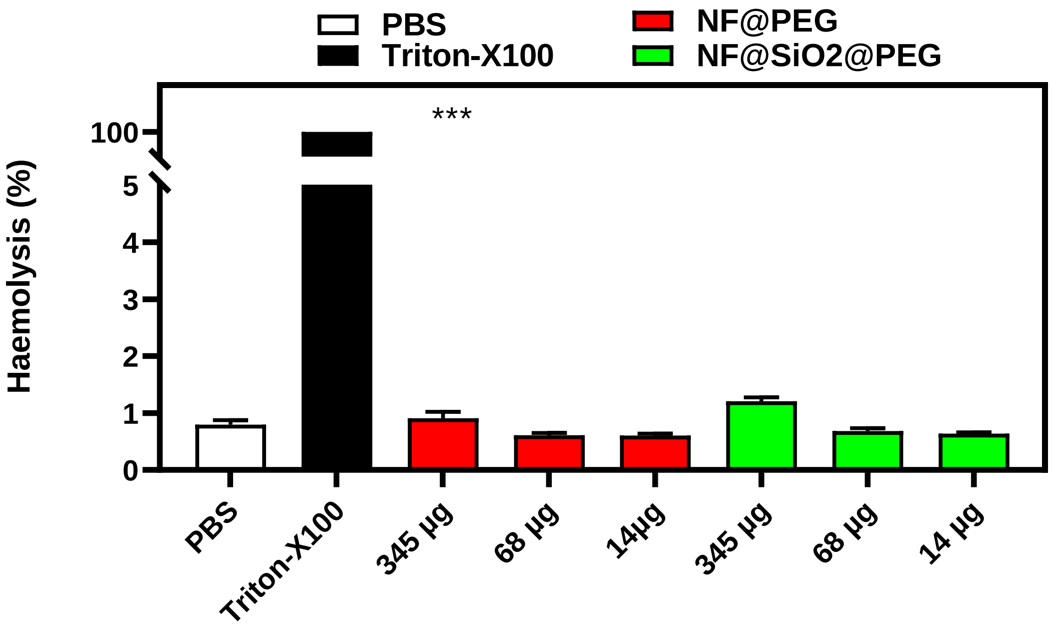

PBS
Triton-X100
NF@PEG
NF@SiO2@PEG
***
Haemolysis (%)
100
5
4
3
2
1
0
PBS
Triton-X100
345 µg
68 µg
14µg
345 µg
68 µg
14 µg


**Figure S7**

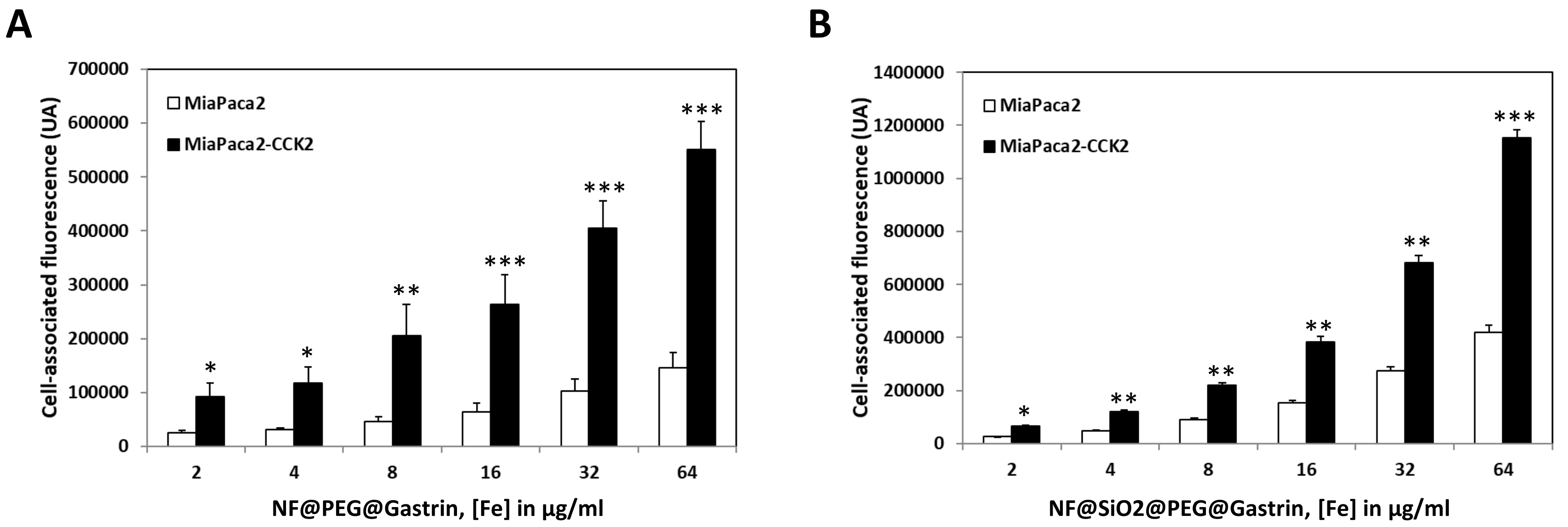
A
700000
600000
500000
400000
300000
200000
100000
0
Cell-associated fluorescence (UA)
MiaPaca2
MiaPaca2-CCK2
*
*
**
***
***
***
2
4
8
16
32
64
NF@PEG@Gastrin, [Fe] in µg/ml
B
1400000
1200000
1000000
800000
600000
400000
200000
0
Cell-associated fluorescence (UA)
MiaPaca2
MiaPaca2-CCK2
*
**
**
**
**
***
2
4
8
16
32
64
NF@SiO2@PEG@Gastrin, [Fe] in µg/ml

**Figure S8**

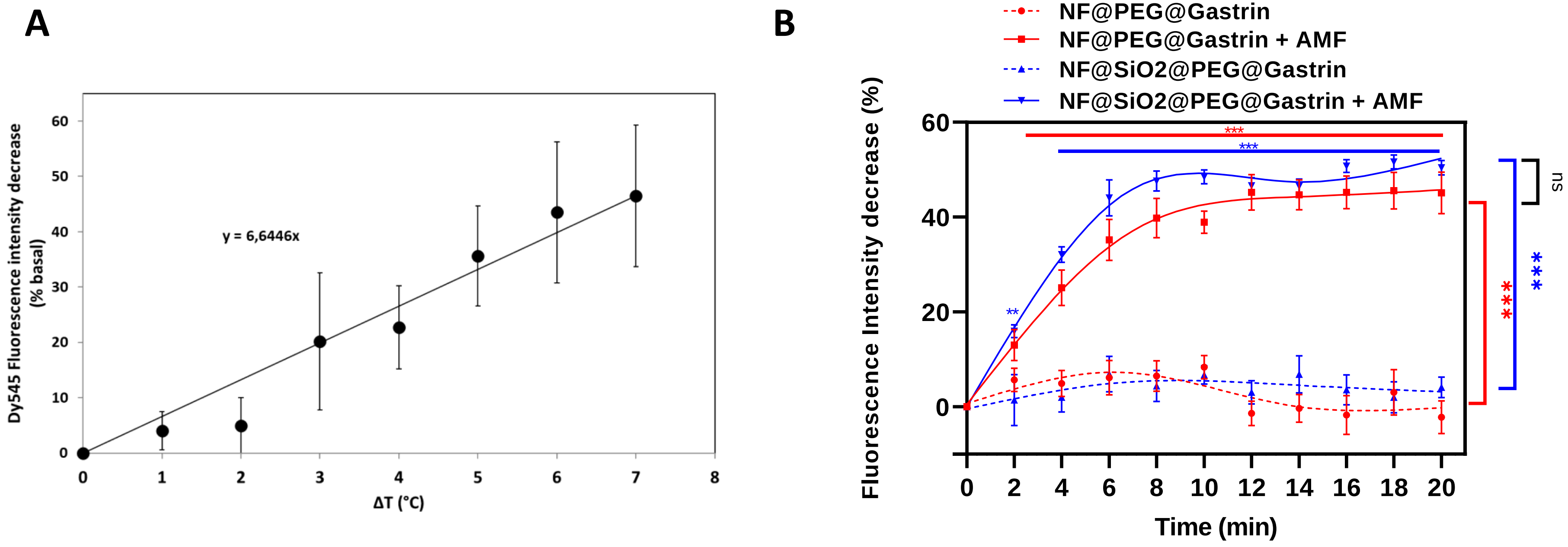
A
Dy545 Fluorescence intensity decrease (% basal)
y = 6,6446x
ΔT (°C)
B
NF@PEG@Gastrin
NF@PEG@Gastrin + AMF
NF@SiO2@PEG@Gastrin
NF@SiO2@PEG@Gastrin + AMF
Fluorescence Intensity decrease (%)
Time (min)
ns
***
**

Figure S9

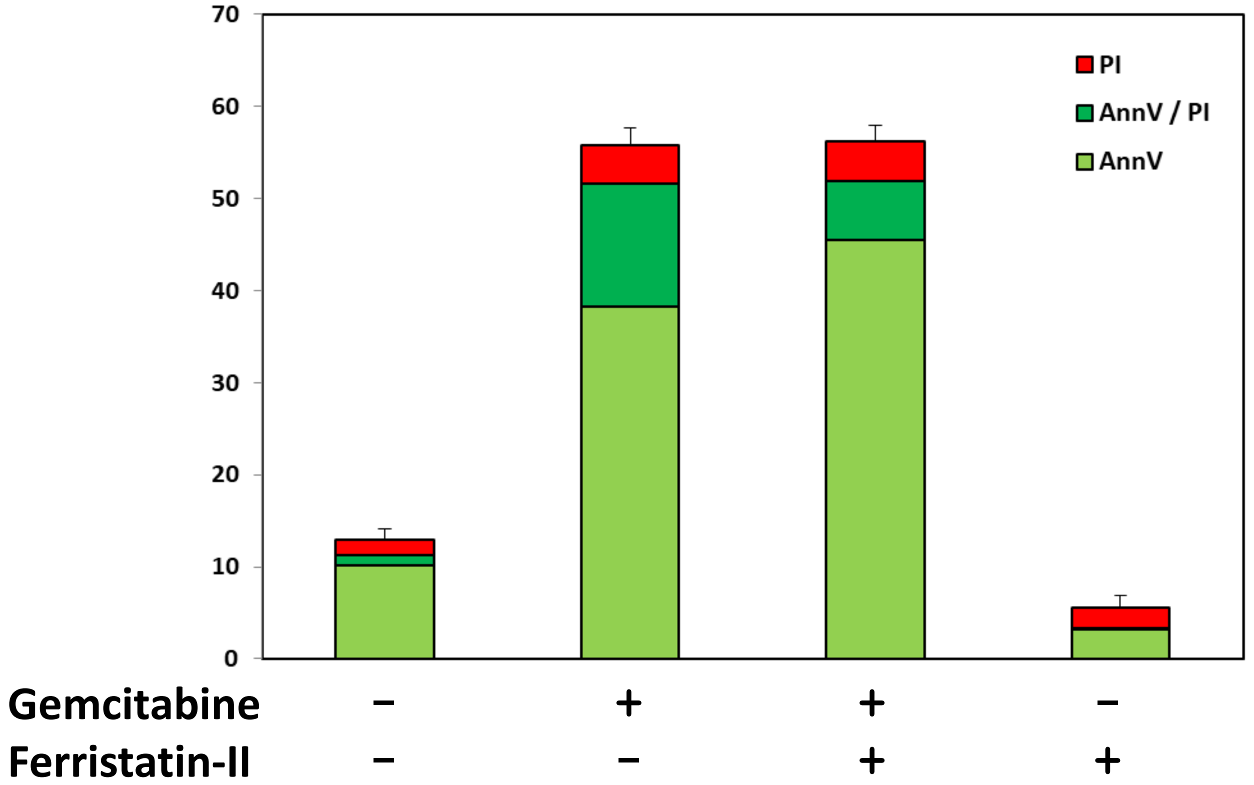
70
60
50
40
30
20
10
0
PI
AnnV / PI
AnnV
Gemcitabine − + + −
Ferristatin-II − − + +

Figure S10
