## Supplementary material for "Origin of Fe ions in ROS production induced in magnetic hyperthermia anti-cancer nanotherapy: release from iron oxide nanoparticles or not?": Legend to supplementary figures

**Figure S1. Simplified illustration of two distinct surface modification techniques to generate unprotected or protected surfaces on γ-$Fe_2O_3$ NFs. (A)** Unprotected coating was done through direct PEG anchoring on bare NFs generating NF@PEG bearing free surface $NH_2$ groups. The latter were used for further coupling with DY647 and Gastrin, necessary to detect and target the IONPs respectively. **(B)** Protective coating was done by firstly growing a silica shell on bare NFs under Stöber conditions. Then, EDPS was added to generate NF@$SiO_2$ IONPs with free -$NH_2$ surface groups, used for covalent PEG-2k-al attachment through reductive amination. The terminal –OH groups of attached PEG were transformed firstly into -Cl groups through $SOCl_2$. Ethylenediamine was then used to generate terminal –$NH_2$ bonds by nucleophile substitution of -Cl groups. The generated $NH_2$ groups on PEG corona were then used to attach DY647 and Gastrin. Each step of surface modification is detailed in Materials and Methods section.

**Figure S2: (A)** TEM image representative of γ-Fe2O3 NF after their synthesis through polyol route and **(B)** its corresponding histogram plotted after manual size determination of N=300 NF using ImageJ software and fitted with a normal distribution. **(C)** TEM image representing the core-shell structure of NF1@$SiO_2$ with and **(B)** its corresponding histogram plotted after manual size determination of N=300 NF@SiO2 using ImageJ software and fitted with a normal distribution.

**Figure S3 1:** AC magnetometry of NF@PEG (A) and NF@$SiO_2$@PEG (B). Hysteresis loop (Mass magnetization vs field intensity *H*) measured at 280 kHz and indicated magnetic field amplitudes.

**Table 1: Physicochemical properties of IONPs.** NF@PEG@Gastrin and NF@$SiO_2$@PEG@Gastrin correspond to NF@PEG and NF@$SiO_2$@PEG nanoparticles functionalized with Dy-647 fluorophore and Gastrin peptide to detect and target them respectively. $D_{TEM}$ value corresponds to the outer diameter of bare NF or NF@$SiO_2$, determined through ImageJ analysis of *N*=300 nanoparticles on TEM micrographs. Hydrodynamic diameter $D_H$ and polydispersity index PDI of IONPs dispersion (0.1 mg γ-$Fe_2O_3$/mL) are measured through DLS in $H_2O$ and calculated through Cumulant method. ζ-potential of IONPs dispersion (0.1 mg γ-$Fe_2O_3$/mL) was measured in 1 mM HEPES with pH set at 7.4. The conformational regime of PEG is given from the ratio between Flory radius RF ($RF$ = $a$(DP)3/5, where a is the effective monomer PEG length = 0.358 nm, DP the degree of polymerization) and average distance D ($D$=2($\pi\sigma$)−1/2) between adjacent PEG chains.

**Figure S4: (A)** Temperature monitoring of IONPs solutions during AMF exposure. **(B,C)** *In vitro* Fe dissolution from IONPs. IONPs solution at 0.5 g/L (B) or 3 g/L (C) in Artificial Lysosomal Fluid (ALF) were incubated in water bath at 37°C or 45°C (corresponding to IONP solution temperature during AMF exposure) or exposed to AMF during 2h. **(D, E, F)** *In vitro* Fe dissolution from IONPs. IONPs solution at 0.5 g/L (D), 3 g/L (E)or 7 g/L (F) in Artificial Lysosomal Fluid (ALF) were exposed to AMF for 2 h. Fe dissolution was analyzed in the surpernatant by UV-VIS spectroscopy.

**Figure S5: (A)** *In vitro* Fe dissolution from IONPs. IONPs solution at 7 g/L γ-$Fe_2O_3$ (B) in Artificial Lysosomal Fluid (ALF) were exposed to AMF during 1 or 2h. γ-$Fe_2O_3$ core diameter were then analyzed by TEM (~300 nanoparticles per condition). **(B)** TEM images of NF@PEG and NF@$SiO_2$@PEG at 7 g/L γ-$Fe_2O_3$: (a) and (d) prior to AMF treatment; (b) and (e) after 1 h AMF exposure; (c) and (f) after 2 h AMF exposure.

**Figure S6: Cytotoxicity of NF@PEG@Gastrin (A) and NF@SiO2@PEG@Gastrin (B) MNPs.** MiaPaca2 and MiaPaca2-CCK2 were incubated with increased concentrations of MNPs during 24, 48 or 72h. Cell viability was determined by MTT assay.

**Figure S7:** Analysis of hemocompatibility of NF@PEG and NF@SiO2@PEG. Murine, rat or human red blood cells from were incubated with increasing concentrations of NF@PEG and NF@SiO2@PEG, 1 mg/ml Triton X- 100 (positive control) or PBS (negative control) for 4h at 37C. Optical density of the supernatant was then measured at 540 nm and the % of hemolysis was calculated relative to the positive control (Triton X-100) inducing 100% hemolysis. Results are expressed as mean ± sem.

**Figure S8: Targeting efficacy of NF@PEG@Gastrin (A) and NF@SiO2@PEG@Gastrin (B) MNPs.** MiaPaca2 and MiaPaca2-CCK2 were incubated with increased concentrations of MNPs during 72h. Cell-associated fluorescence was measured by flow cytometry, and results are expressed as fluorescence associated with the cells and are the mean ± SEM of at least four separate experiments.

**Figure S9: Fluorescence intensity of NF@PEG@Gastrin or NF@SiO2@PEG@Gastrin is dependent on temperature.** MiaPaca2-CCK2 cells were incubated with Gastrin-MNP-DY549 for 72h. **(A)** Cells were slowly heated in the incubator chamber of the confocal microscope. Fluorescence intensity of fluorophores was analyzed from confocal microscopy images, normalized to $\Delta T=0$ and expressed as the mean ± SEM of at least 4 separate experiments. **(B)** AMF was applied using a miniaturized electromagnet. Temperature at MNPs surface was analyzed by measuring the decrease of theDY549 fluorescence intensity. 20–30 cells/experiments were analyzed, and results are the mean ± SEM. Significant differences comparatively to t = 0 are indicated above the curve; significant differences between conditions are indicated with brackets.

**Figure S10: Analysis of cell death induced by gemcitabine in presence or not of Ferr-II.**
Cells were incubated or not with gemcitabine (10 µM) for 72h, and with Ferr-II in DMEM medium without SVF for 24h. Dead cells were labeled with AnnV/PI and counted 4 h after AMF exposure by confocal microscopy. Results are the mean ± SEM of 4 separate experiments.
